# Agent-based simulations of lung tumour evolution suggest that ongoing cell competition drives realistic clonal expansions

**DOI:** 10.1101/2025.11.04.686578

**Authors:** Helena Coggan, James R. M. Black, Carlos Martínez-Ruiz, Kristiana Grigoriadis, Jasmin Fisher, Nicholas McGranahan

## Abstract

Computational simulations of tumour evolution are increasingly used to infer the rules underlying cancer growth, with the goal of one day recommending tailored treatments. To make reliable inferences, such models must be able to reflect the properties of real tumours. Recent work has shown that lung tumours undergo frequent and late subclonal expansions, which are associated with poor prognosis. This paper tests three candidate simulations of three-dimensional tumour growth, which make different assumptions about the nature of competition between cells, for their ability to replicate these late expansions. Only a model which assumes stringent competition between tumour cells for existing tissue space, after the tumour has reached a fixed size, can produce multiregion sequencing data realistic to lung tumours. This work also assesses the influence of these assumptions on the inferred selection strength of individual tumours in a large cohort of lung cancers, and finds that inferences from a well-tailored model produce much higher estimates than poorly-tailored models of the effects of driver mutations on cell fitness.

**Table of Contents:** Computational simulations of tumour evolution are increasingly used to infer the rules underlying cancer growth, with the goal of one day recommending tailored treatments. Here we show that the properties of lung cancer sequencing data are best replicated by a model which assumes that cells compete both to proliferate and survive.

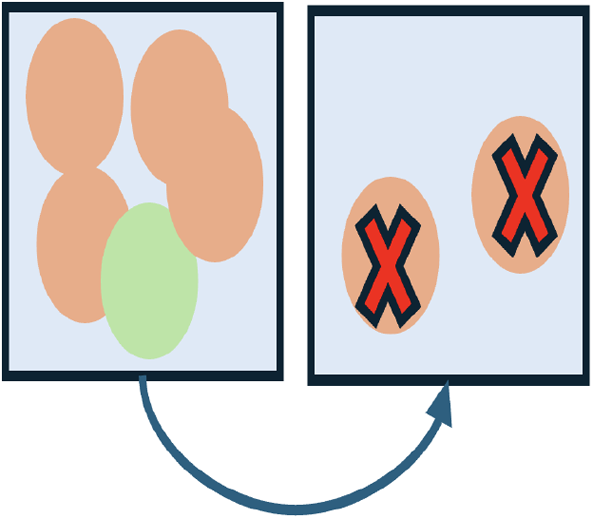

## 1 Introduction

In the last decade, spatial agent-based models (SABMs) have become an increasingly popular tool for investigating the underlying rules of cancer evolution [1, 2]. Such models usually start with a single cell in the center of a two-dimensional [3, 4, 5, 6, 7, 8, 9] or three-dimensional grid [10, 11, 12, 13, 14], and simulate cell division, death and acquisition of mutations until the tumour is sufficiently large. At this point, the simulated tumour is sampled, and these samples are sequenced, to measure the number and frequency of the mutations present. These simulated data can then be compared to real tumour samples. In this way, SABMs can be used to infer the evolutionary rules driving the growth of specific tumours.

This approach leverages the increasing availability of multi-region sequencing (MRS) tumour data, which provide a clearer picture of the evolutionary trajectory of a tumour by measuring the distribution of mutations across different regions and quantifying the ‘relatedness’ of different parts of the tumour [8, 11]. By developing simulations to decipher the site-specific rules of tumorigenesis, researchers can develop a deeper understanding of the rules and modes of tumour development, which may ultimately inform strategies to prevent and treat cancers.

To produce reliable inferences, however, the model must be able to reflect the biology of real tumours. Historically, SABMs have generally been modelled either on colorectal cancer (CRC) [15, 16, 17, 10, 11, 8], due to the relative abundance of MRS datasets in CRCs, or ‘general’ solid tumours [18, 6, 12, 14, 19]. To our knowledge, no SABM of cancer has included realistic local competition for survival, representing the idea that a cell is less likely to survive when surrounded by a large number of fitter cells (i.e. ‘active competition’ [20]). This is a significant limitation when attempting to infer selection strength (i.e. the fitness advantage conferred by beneficial mutations) across cancer types, because it encodes an assumption about how selection ‘works’ that may not apply to all tumours. This is especially true of lung tumours, which have patterns of intra-tumoural heterogeneity suggestive of large, frequent regional expansions by ‘subclones’ of related cells (illustrated in Figure 1). These regional subclonal expansions have been linked to poorer prognosis [21]. To the best of our knowledge, no modelling framework exists which has been specifically developed for, or evaluated against, the evolutionary patterns of lung tumours. As a result, there is an unmet need for a model which can reliably infer the evolutionary rules of individual lung tumours, allow researchers to assess the extent to which these rules vary between patient’s tumours, and to understand the extent to which this may influence treatment outcomes.

**Figure 1.**
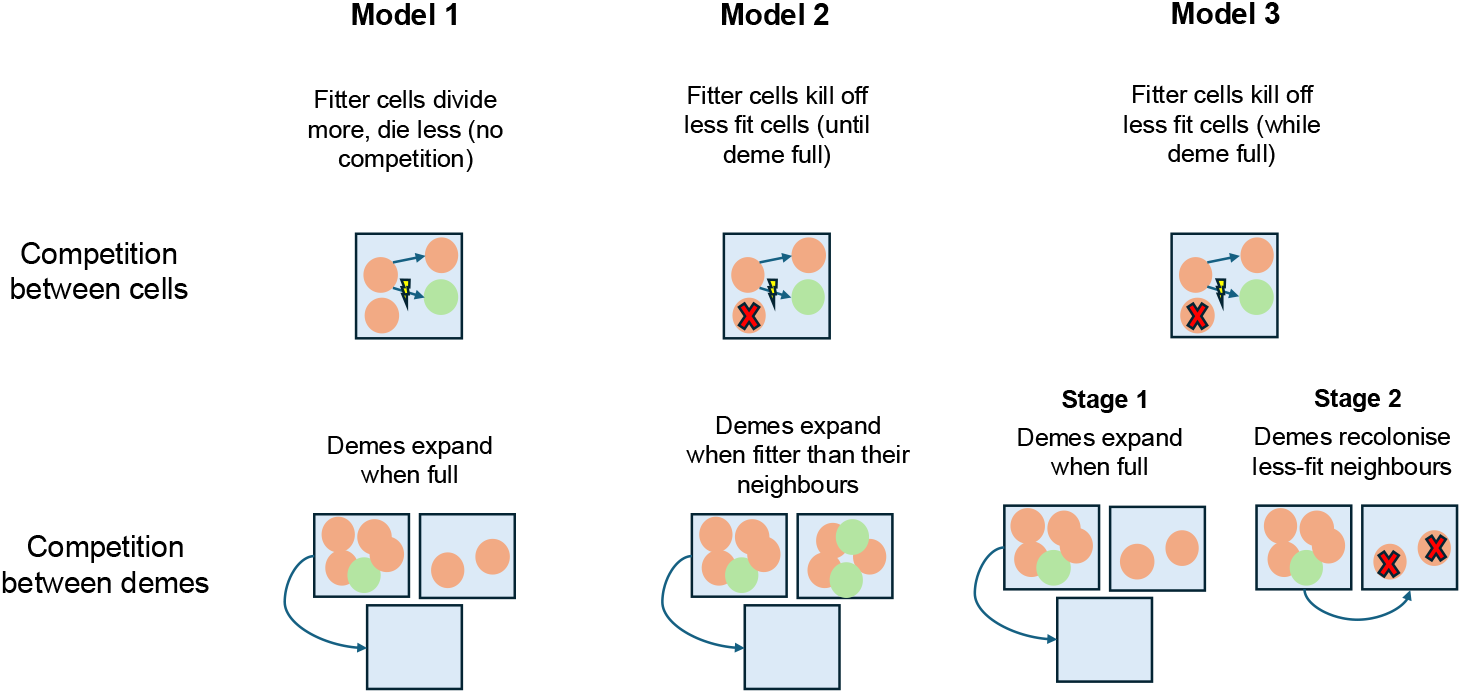
A schematic of the regional subclonal expansions characteristic of lung tumours in the TRACERx cohort. All cells in the tumour are descended from a single cell, the most recent common ancestor (MRCA, marked in grey). At the end of the tumour’s growth, it is resected and subject to multi-region sequencing. During tumour growth, certain lineages spread and fix in regions of the tumour without ‘sweeping’ the entire lesion and becoming the MRCA. Mutations present in the cell marked in orange are detected in Sample 1, fixed (detected in all cells) in Sample 2 and absent from Samples 3 and 4. Mutations present in the cell marked in green are fixed and private in Sample 4.

In this paper, we present three agent-based models of tumour growth, each encoding different assumptions about selection. We compare simulated multi-region sequencing data from all three models to properties of real lung tumours, drawn from publicly available MRS data from a cohort of treatment-naive lung tumours (the TRACERx dataset [21]). These models represent increasingly stringent forms of cell competition, at the cost (given finite computational resources) of decreasing final tumour size (see Section 4). We vary the selection strength and death rate within each model, and find that only the third model, in which cells compete for survival both within and between demes, is capable of replicating the late, region-specific clonal expansions characteristic of NSCLC tumours in the TRACERx cohort (Section 2).

We also examine how accurately selection can be inferred from simulated data, leveraging methods from machine learning and Approximate Bayesian Computation (Section 2.4), and demonstrate that model choice has a profound impact on the selection strength inferred from TRACERx tumours. In particular, an inference process which relies on a model which is incapable of replicating region-specific clonal expansions will tend to significantly underestimate the selection strength present in lung tumours, thus potentially leading to erroneous conclusions regarding the extent and importance of positive selection.

This work defines a new ‘state of the art’ for lung tumour modelling, and emphasises the importance of developing tumour-specific models to infer the rules of cancer evolution. Each model can be run in Python, with code publicly available in a Github repository (hcoggan/Agent-based-simulations-of-lungtumour-evolution).

## 2 Results

### 2.1 Agent-based models can capture local competition for survival and expansion

We present here three candidate models of tumour evolution. All three assume that changes in cell fitness arise as a result of somatic driver mutations of selection strength *s*, acquired when cells divide. All three also assume that cells divide and die within small areas of tissue (‘demes’) of 10,000 cells, and that the tumour expands through three dimensions as cells colonise new demes.

However, each model makes different assumptions about the effect of selection on a cell’s proliferation or survival. In Model 1 (adapted from SCIMET [11]), cells are increasingly likely to divide as they acquire driver mutations, but there is no direct competition for survival, either within or between demes. The two subsequent models introduce increasingly severe competition for space and resources. In Model 2, cells compete for survival within demes as the tumour continues to expand. In Model 3, the tumour expands to a fixed size and then continues to evolve during a second ‘homeostatic’ stage of growth, in which cells compete for survival both within *and* between occupied demes.

Whilst all models make different assumptions about the way selection influences tumour evolution, they share a common set of assumptions about spatial structure, mutation rate and three-dimensional growth dynamics (see Figure 2). We next test whether each model can replicate the key characteristics of lung cancers.

**Figure 2.**
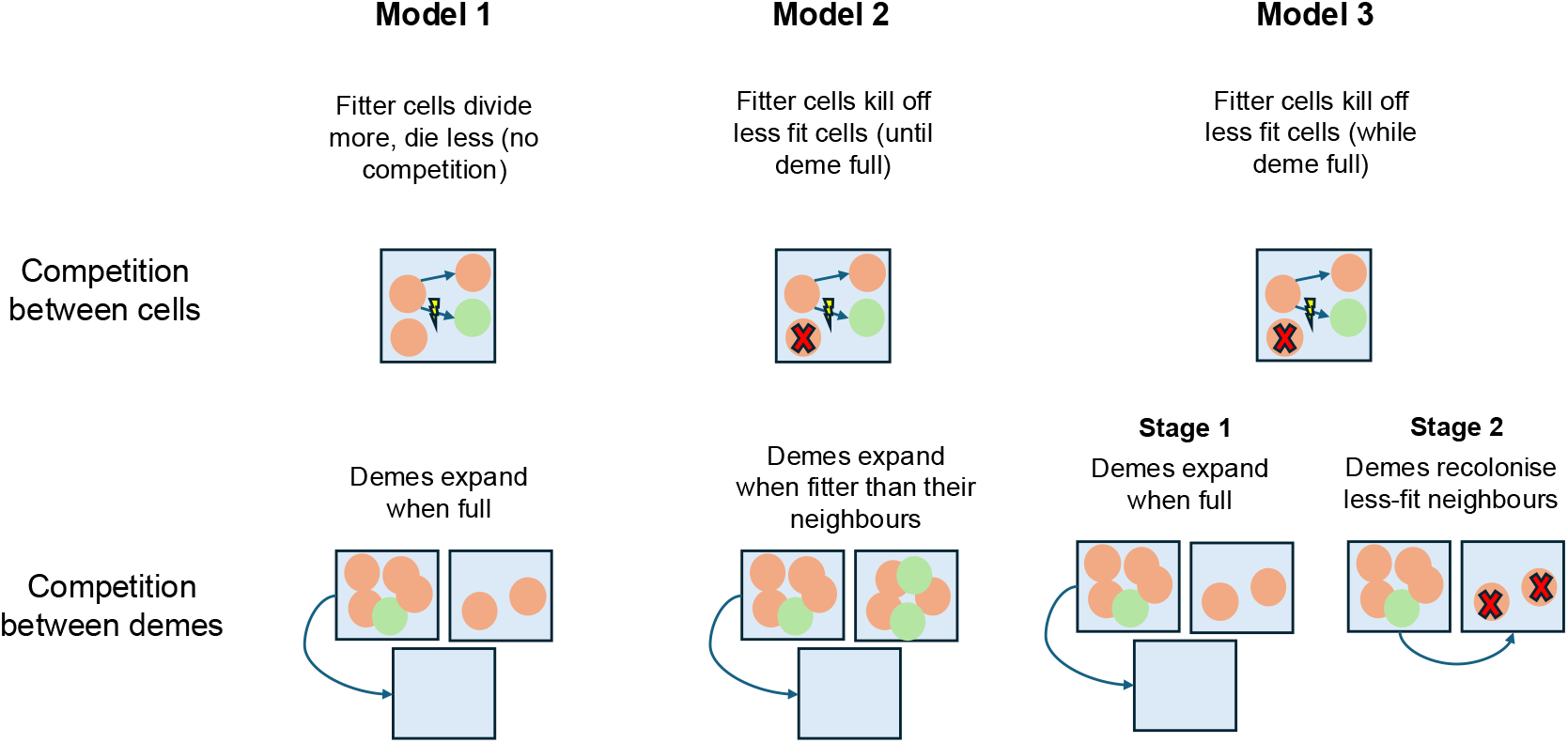
A comparison of the three models evaluated in this paper. ‘Lightning bolts’ indicate the acquisition of a genetic alteration; red crosses indicate cell death; arrows between cells indicate division; arrows between demes indicate the migration of half the cells in one deme to another.

### 2.2 Comparing simulated to real tumours with multi-region sequencing data

We first assess the ability of each model to replicate the properties of lung cancer. We use three metrics to compare simulated tumours to cancers. Firstly, for each summary statistic *S*_*i*_, we can measure the distribution of statistics *S*_*i*_(*s, M*) produced by tumours with selection strength *s*, and *M* observed samples (for Models 1 and 2, we can also vary cell death rate; see Supplementary Materials). We are interested initially in whether the average of *S*_*i*_(*s, M*) is ‘TRACERx-plausible’, i.e. whether it falls within the 90% confidence interval for the corresponding summary statistic in comparable real tumours (i.e., those from which *M* samples were taken). We are also interested in whether we can reproduce the *median* value of that summary statistic in the TRACERx cohort, i.e. whether models can capture the properties of ‘average’ lung tumours and not simply the extremes.

We focus on two key summary statistics: the simulated tumour mutational burden (TMB), the number of mutations observed across the tumour (Figure 3); and the number of clonal illusions (CIs), mutations which are fixed (CCF=1.0) in at least one region but not all (Figure 4).

**Figure 3.**
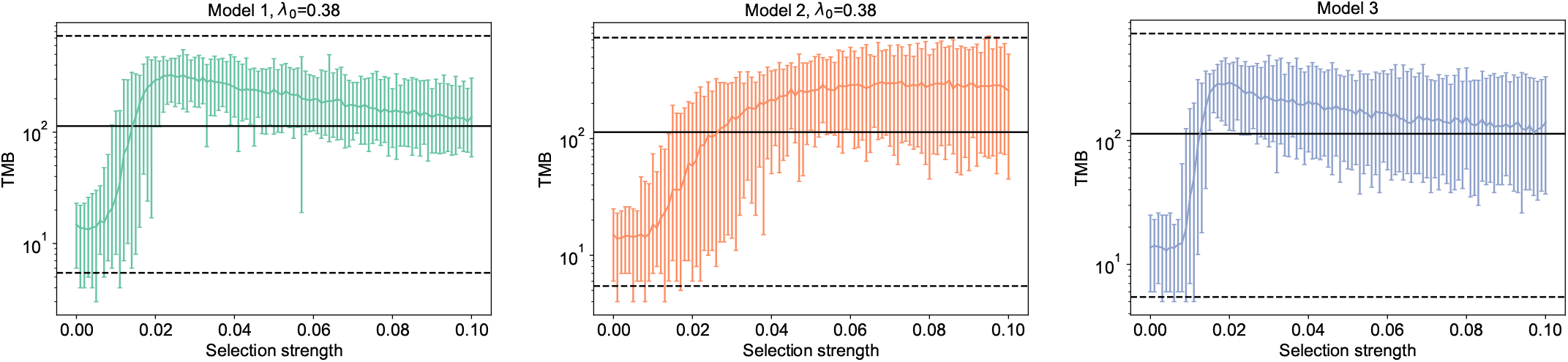
The variation of simulated TMB in each model with selection strength *s*. Models 1 and 2 have an assumed death rate of *λ*_0_ = 0.38. We assume *M* = 2 samples. Averages are taken from 100 repeats at each *s* value, with maxima and minima indicated by error bars. Dotted black lines indicate the outer limits (5th and 95th percentiles) of corresponding summary statistics from all TRACERx tumours which were sampled at *M* regions. Solid black lines indicate the medians of these real TRACERx tumour cohorts.

**Figure 4.**
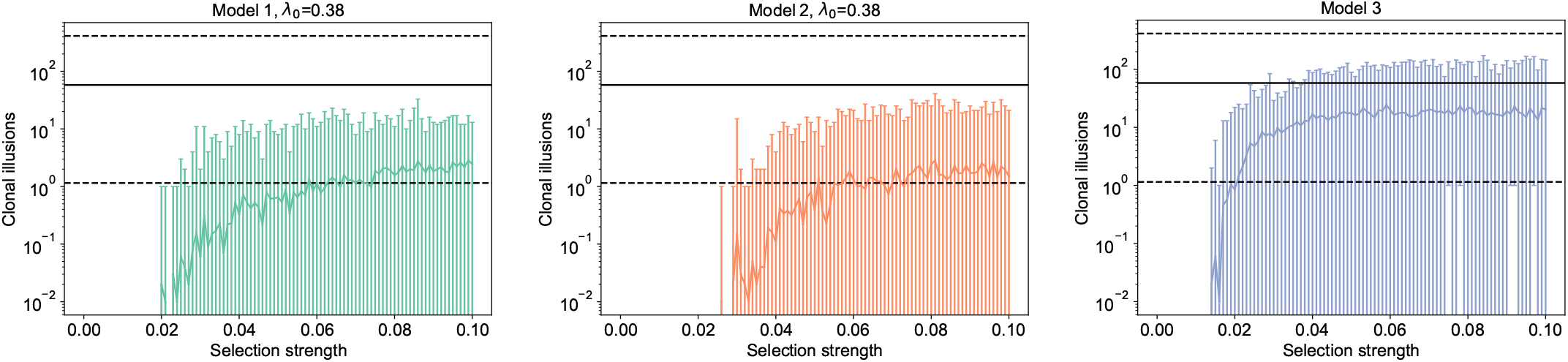
The variation of simulated clonal illusions (CIs) in each model with selection strength *s*. Models 1 and 2 have an assumed death rate of *λ*_0_ = 0.38. We assume *M* = 2 samples. Averages are taken from 100 repeats at each *s* value, with maxima and minima indicated by error bars. Dotted black lines indicate the outer limits (5th and 95th percentiles) of corresponding summary statistics from all TRACERx tumours which were sampled at *M* regions. Solid black lines indicate the medians of these real TRACERx tumour cohorts.

We can also consider *N*_*truncal*_, the number of mutations observed at a CCF of 1.0 in all samples (and thus excluded from the calculation of all other summary statistics, *S*_*i*_). As discussed, the presence of an unknown number of somatic mutations in the founding cells of real tumours makes it impossible to usefully compare *N*_*truncal*_ between real and simulated tumours. However, differences in *N*_*truncal*_ *between* simulated tumours can still be informative about the underlying model dynamics. In particular, in a realistic model of lung cancer, increasing selection strength should allow later-arising lineages to dominate the tumour, and so extend the latest possible point at which a driver mutation can arise and still become the ancestor of all sampled cells. Higher selection should drive more and later clonal sweeps, such that the MRCA has more mutations relative to the founding cell; thus *N*_*truncal*_ should increase with *s*. We examine this in Figure 5.

**Figure 5.**
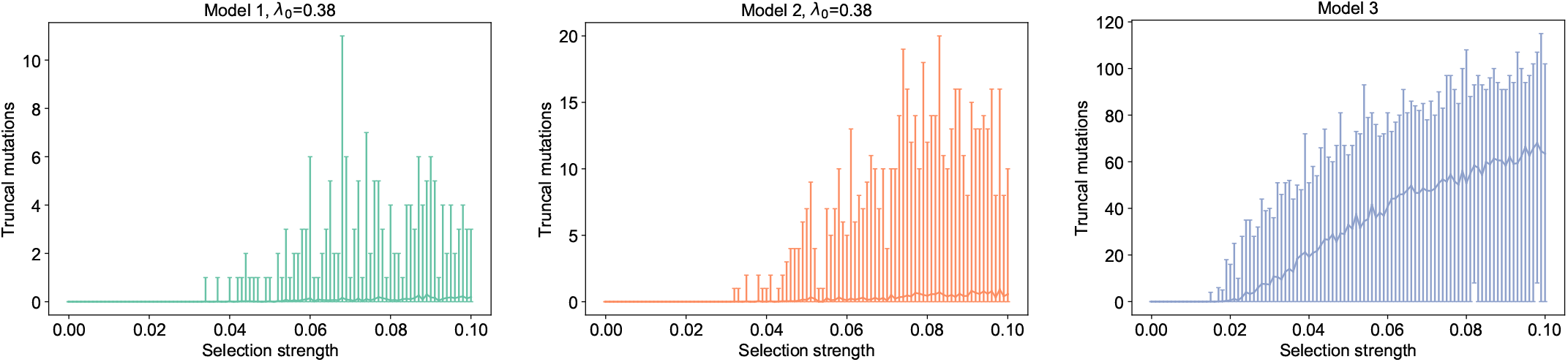
The variation of the number of truncal mutations, i.e. those detected at a CCF=1.0 in every region (and are thus excluded from the calculation of all other summary statistics) in each model with selection strength *s*. Models 1 and 2 have an assumed death rate of *λ*_0_ = 0.38. We assume *M* = 2 samples.

#### 2.2.1 Intra-tumoural heterogeneity is greatest at intermediate levels of selection

We first observe that for models with a variable death rate (Models 1 and 2), realistic TMBs are generally only achieved for the highest possible death rate (*λ*_0_ = 0.38, corresponding to growth rates of 1% per day). This would imply that tumours have a final age of around 5 years; previous estimates have suggested that both lung adenocarcinomas (LUADs) and lung squamous cell carcinomas (LUSC) are diagnosed roughly 2 years after the development of the most recent common ancestor (MRCA) [22], an event which occurs within the timeframe of our model (which begins post-transoformation).

Even where selection is strong, TMBs remain resolutely low for death rates below the maximum at *λ*_0_ = 0.38 (see Supplementary Figures 8 [Model 1] and 9 [Model 2]). This dependence on death rate is fairly intuitive: an increased *λ*_0_ results in higher cell turnover, which increases the time taken for tumours to reach full size and thus the mitotic age of the sampled cells. Cells acquire mutations when they divide, so this higher death rate leads to a higher TMB. A higher baseline death rate also increases the selective survival benefit (i.e. decrease in death rate) of driver mutations, and so will accelerate the pace of evolution.

Once we fix *λ*_0_ = 0.38 for Models 1 and 2, all models can produce an equally realistic range of TMBs (see Figure 3), independently of sample number (see Supplementary Figures 5 and 6. This is to say that different assumptions about competition do not strongly affect our ability to produce *diverse* tumours (using TMB as a measure of diversity).

The relationship between TMB and selection strength can also inform our understanding of the emergent growth dynamics in each model. In Models 1 and 3, cells either do not compete directly for survival (1) or compete only to recolonise existing tissue areas (3), but do not compete explicitly for new space. In both models, we observe that selection strength first increases and then decreases the mean TMB. First, we observe a peak TMB at weak selection (*s* ≈0.02) (see Figure 3, left panel). For lower selection levels (*s* < 0.01), competition is weak, and so very few mutations reach detectable prevalences; those which do are likely to have occurred early, and thus appear in multiple samples, such that the resulting tumour is very homogenous. As competition increases, the sampled population begins to diversify. Selection is now strong enough to allow lineages with late-occurring driver mutations to increase in frequency: sampled populations are composed of a large number of highly proliferative lineages, frequent enough to be detectable but not sufficiently advantaged to take over an entire region.

As selection strength increases further (*s* > 0.02), the TMB begins to decrease again. Selective advantages are now strong enough that fitter lineages proliferate very quickly. These lineages invade empty space aggressively enough to block less-fit lineages from reaching the surface. If this process occurs sufficiently early, these cells become the ancestors of multiple samples, increasing the genetic relatedness of sampled regions. This makes it less likely that any sampled mutation will be private, and decreases the overall number of genetic mutations (TMB) contained in the sampled regions. Similar patterns are observed in the number of mutations observed only in 1 tumour region (TMB-priv, see Supplementary Material).

In Model 2, however, selection strength has a saturating effect on the TMB. Tumour diversity is very low at *s* = 0.0, increases slowly as selection increases, and then plateaus after *s* = 0.05. This suggests that explicit competition for new space introduces a ‘bottleneck’ on selection: beyond *s* = 0.05, selection strength does not increase the diversity of the tumour. This in turn implies that there is a point in tumour growth at which it is simply too late for any new lineage, however fit, to reach detectable prevalence. This limits the ability of Model 2, which assumes that cells compete for expansion into new tissue but not into already-colonised tissue, to replicate the competitive dynamics of lung cancer.

#### 2.2.2 Competition for existing space is necessary to simulate late subclonal expansions

We define a clonal illusion as a mutation which is fixed in *at least one region*, but is subclonal throughout the entire tumour. As all truncal mutations are excluded from our main summary statistics, this latter condition is automatically satisfied. The number of clonal illusions in a tumour with *M* samples is therefore

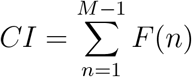

where *F* (*n*) is the number of mutations fixed in exactly *n* samples. If no CIs (or almost none) are observed, we can say that TRACERx-like subclonal expansions do not occur in the model.

We observe that Models 1 and 2 are unable to replicate realistic numbers of clonal illusions at any selection level (see Figure 4). CIs are almost non-existent for *s* < 0.05 (i.e. with a mean of fewer than one per simulated tumour) and essentially constant for *s* ≥ 0.05. This pattern is independent of the number of sampled regions *M* (Supplementary Figures 12 and 13).

These results suggest that indirect competition (as in Model 1) or competition for new space alone (as in Model 2) are insufficient to allow late-arising, fit lineages to take over individual regions before the tumour is sampled. On average, these models produce fewer than 10 clonal illusions per tumour, regardless of sample number. This is true even at high selection levels, where the TMB and TMB-priv *are* in line with those of observed tumours. This suggests that we could not obtain competitive evolutionary dynamics under this model, even if it were computationally feasible to simulate larger tumours: the speed at which mutations spread is simply too slow relative to the rate at which the tumour diversifies. This leads us to a two-stage model of growth in which cells compete for *existing* space after an initial expansion (Model 3).

Model 3 produces realistic levels of clonal illusions at high selection levels (*s* > 0.02) (see Figure 4, bottom panel). As with the TMB (Figure 3), the number of clonal illusions continually increases with ongoing selection. We infer from this that mutations arising later in tumour evolution can spread beyond their sample of origin, so functional competition is ongoing and each increment of selection provides a continuing advantage. Continuing evolution allows lineages to spread through regions in a fixed-size tumour.

Simulations are more likely to produce sufficient levels of CIs at low sample number (see Supplementary Figures 12 and 13, bottom panels), and the range produced is still at the lower end of the observed data (on average fewer than 100, regardless of sample number, when hundreds would be permissible). The *average* number of simulated CIs, whilst within the 90% confidence interval for the TRACERx dataset, is always lower than the median for real tumours; realistic values are produced only by ‘extreme’ simulations. Model 3 nonetheless represents a high-water mark for our ability to replicate the properties of real lung cancers. This in turn suggests that the assumption of ongoing competition for existing space allows us to accurately model the underlying biology. We can confirm this by examining the relationship between selection and ‘clonal sweeps’.

#### 2.2.3 Clonal sweeps rely on ongoing competition throughout tumour growth

We finally consider the relationship between selection strength *s* and *N*_*truncal*_, the number of mutations fixed in all *M* samples and thus excluded from our main summary statistics. As discussed, these cannot be directly compared to the TRACERx cohort, as truncal mutations in real tumours may have occurred pre-transformation (i.e. before the start of the timeframe considered by the simulations). However, patterns of truncal mutations can illustrate the emergent dynamics of simulated tumour growth.

In Model 1 (Figure 5, topmost panel) we observe that the number of truncal mutations is extremely low, fewer than 10 in all cases and no higher than 1 when more samples are taken (see Supplementary Figures 14 and 15, topmost panels). This strongly implies that the founding cell of the tumour is generally the MRCA of all sampled tumour cells, and that no clonal sweeps occur after tumour initiation.

An exome mutation rate of *µ* = 0.4 per cell division suggests that, in cases where only one mutation is clonal (meaning that the MRCA only has one more mutation than the founding cell), that mutation appeared within a few divisions of tumour initiation; if clonal sweeps occurred later, then the MRCA would likely have accumulated more mutations during its own evolution, and those mutations would also have become clonal. Model 1 therefore produces ‘branching growth’, i.e. unbroken diversification from the founding cell.

We can confirm this by examining the number of mutations observed in all regions (see Supplementary Figure 14), which increases with selection. Combined with the absence of clonal illusions (see the top panel of Figure 4 in the previous section), we may conclude that selection drives the emergence of fitter lineages which maintain a ‘universal’ subclonal presence but rarely take over entire regions. In general, competition between cells in Model 1 is not sufficient to allow clonal sweeps or regional subclonal expansions: fitter lineages slowly increase in prevalence as the tumour grows, but never eliminate their less-fit competitors. These growth patterns are laid out in Figure 6 (top panel).

**Figure 6.**
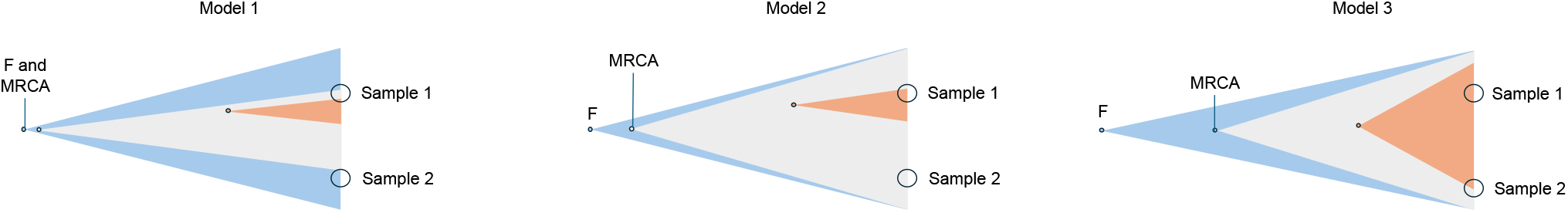
A fish plot of the growth patterns resulting from the indirect-competition model (Model 1, left), the directcompetition model (Model 2, middle) and the two-stage direct competition model (Model 3, right). The grey and red clones are successively fitter than the founding lineage (blue clone). In Model 1, the grey clone increases in frequency but never fully eliminates the blue clone, and is found subclonally in both samples. The red clone appears later during tumour growth and is found subclonally in one sample. In Model 2, the grey clone arises early enough to eliminate the blue clone, becomes the MRCA of the tumour, and is observed clonally in both regions. The red clone, though fitter, arises too late to seed both samples and is observed (clonally or subclonally) in Sample 1. In Model 3, the red clone arises late in tumour growth and is able to take over Sample 1 (so is detected clonally there) and spread to Sample 2 (so is detected subclonally).

Model 2, however, successfully produces tumours with clonal sweeps (see 5, middle panel). Unlike Model 1, in which no tumour ever has more than 1 truncal mutation if sufficiently sampled, the inclusion of direct competition in Model 2 produces simulated tumours with 10-20 truncal mutations (This decreases as more regions are sampled: see Supplementary Figures 12 and 13, middle panel.) This suggests that fit lineages which appear early enough in lesion growth– within the first few dozen cell divisions after transformation– can drive competing lineages to extinction and become the MRCA of the entire tumour. After this, no mutation is able to fix across the entire tumour, even when it is left to grow to ∼10^8^ cells (and ∼10^4^ demes); all later mutations are sample-private. We also observe that in Model 2 selection does not increase the number of mutations seen in all regions (see Supplementary Figure 16, middle column).

Taken together, these observations suggest that direct competition for expansion in Model 2 in fact *limits* the spread of fitter lineages, relative to Model 1 (no competition). Model 2 assumes that the tumour expands into normal tissue in ‘waves’ when its total pressure becomes too high, with surface cells competing with each other for newly available space and resources, then this limits the speed with which mutations can spread through the tumour. A late-arising lineage, even if it quickly outcompetes all other cells in its deme, will not be able to invade multiple sampled regions if it arises when the tumour is almost grown and has only a few ‘expansions’ left. Almost all relevant competition occurs early, when there is still time for a lineage which outcompetes its neighbours to become the ancestor of some or all of the sampled tumour regions. This constraint acts as a ‘bottleneck’ on the effect of selection: a selection coefficient of *s* = 0.1 is no more likely to result in a clonal sweep (see Figure 5) or clonal illusion (Figure 4, middle column) than *s* = 0.05. We can conclude that in Model 2, growth is dominated by a small number of mutations which sweep the tumour when it is very small, after which lineages compete within samples but rarely escape their region of origin (see Figure 6, middle column).

We observe that Model 3, which assumes that cells can recolonise existing tissue, *can* produce repeated clonal sweeps. The average number of truncal mutations, *N*_*truncal*_, is an order of magnitude higher than Models 1 and 2 (averages can reach up to 60 at high selection levels, when all previous averages were lower than 2), and increases almost linearly with selection after *s* = 0.02 (see Figure 5, bottom panel). The lack of a saturation curve suggests that competition for existing space (Model 3) removes the bottleneck introduced by direct competition for new space (Model 2): there is no longer a hard ‘time limit’ by which a fit lineage must emerge to fix in multiple samples. We conclude that the introduction of direct competition for *existing* space in Model 3 allows the continual replacement of less-fit clones with fitter cells. We therefore observe the late regional expansions characteristic of real lung cancers (see Figure 6, bottom panel).

### 2.3 Competition for existing space provides the best approximation to the properties of lung tumours

The measurements above suggest that, whilst computationally expensive, competition for existing space as in Model 3 (rather than for new space in an expanding tumour, as in Models 1 and 2) is necessary to accurately replicate the dynamics of ongoing subclonal expansion in lung cancer. A full comparison of summary statistics in all three models for *M* = 2 is provided in Figure 7 (see Supplementary Figure 17 and 18 for *M* = 5, 8). We find that, across all summary statistics, tumours from Model 3 compare better to the TRACERx cohort than those from Models 1 and 2, regardless of death rate.

**Figure 7.**
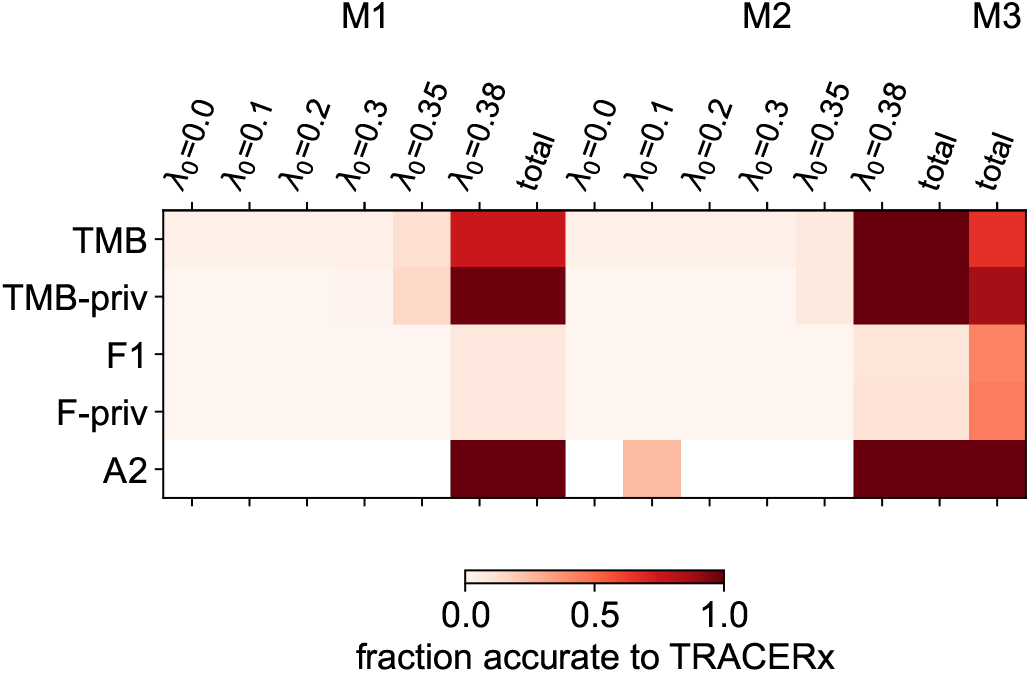
A comparison of the suitability of all 3 models to the TRACERx data, assuming *M* = 2 samples. The element in the *i*th row and *j*th column indicates the probability that a simulation from model *j* will produce a summary statistic *i* within the 90% confidence interval of TRACERx tumours with *M* samples. Models 1 and 2 are broken out by death rate; the ‘total’ column displays the average across all death rates.

**Figure 8.**
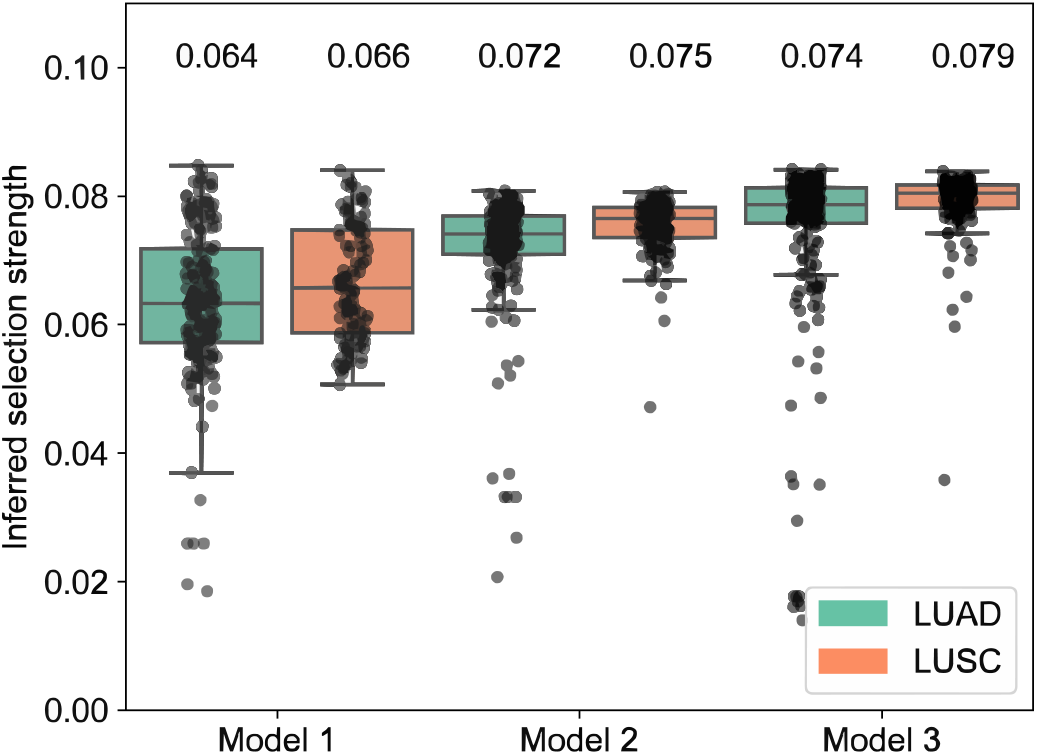
The selection strength inferred from each tumour in the TRACERx cohort using each model as a baseline. Each dot represents one tumour. For Model 1 and Model 2, we assume the maximum death rate of *λ*_0_ = 0.38. Predictions are adjusted for sample number: selection is inferred from tumours with *M* samples using an ABC-ML pipeline trained on simulated tumours with *M* samples. Inferred selection values are averaged across 10 repeats, where each repeat uses a model trained on a different train-test split of the simulated tumours. Right: the mean standard deviation 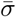 of the posterior distributions inferred from each of 10 repeats, defined as 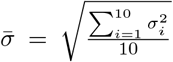. A higher standard deviation suggests greater uncertainty.

We note that ‘fixed and private’ mutations are systematically underestimated amongst all models and all numbers of sampled regions (though Model 3 remains the best at replicating these subclonal expansions). This attribute cannot be easily adjusted using model parameters, because changes which make it easier for a mutation to become fixed in a region (e.g. an increase in selection) will also make it easier to spread beyond that region, which is to say it will make it harder for mutations to stay *private*. It is also likely that the detection thresholds we apply to simulated mutations, as described in Section 4.3, do not fully reflect the complexity of the filters applied to real TRACERx tumours. These depend on aspects of lung tumour biology not explicitly included here (copy number alterations, tumour purity, and so forth), and may be more conservative when determining whether a mutation is detected in multiple regions (and so more likely to classify it as ‘private’). It is also possible that tumour and effective deme size (i.e. the size of the neighbourhood within which cells compete locally) vary between tumours, and that tumours with more fixed-and-private mutations tend to be larger or have smaller effective deme sizes. Our assumption that all tumours have a fixed ‘deme size’ may be limiting our ability to replicate subclonal expansions, even as it increases the computational feasibility of the study.

Whilst we cannot currently capture the numbers of clonal illusions seen in all tumours, Model 3 nonetheless represents a significant advancement in three-dimensional models of solid tumour evolution– specifically in its ability to replicate the regional subclonal expansions seen in NSCLC. We can now leverage this new, more accurate model to infer the selection strength of individual tumours in the TRACERx cohort.

### 2.4 Assessing selection strength in tumours using Bayesian inference

Two- and three-dimensional model frameworks are frequently used to make tumour-specific inferences about cancer evolution (e.g. tumour age, time of metastasis seeding, fraction of cells which are stem cells, etc). We have seen in Section 2 that model assumptions about cell competition– whether competition is direct or indirect, for new or existing space, and so on– can have profound effects on the simulated growth dynamics. These differences may have knock-on effects on the results of inferences from real tumours. Here, we test this proposition by using the three models discussed above to infer selection strength from specific tumours in the TRACERx cohort.

Our procedure for inferring continuous selection strength *s* from tumour data incorporates both ABC and ML. Here we take half of the cohort of simulated tumours and train a regressor (XGBoost-Regressor) to predict *s* from their summary statistics. The difference between the predictions assigned to two tumours then becomes our distance function to assess the dissimilarity of those tumours. We use the other half of the simulation cohort to perform ‘traditional’ ABC (as used by Hu and coauthors [11]) using this distance function. Our strategy amounts to constructing a probability distribution for selection strength *s* from the properties of the simulated tumours which a regressor ‘believes’ to be most similar to the real tumour (see Supplementary Methods for details). We test this pipeline on simulated tumours and find that the choice of model does not affect the pipeline’s ability to determine selection strength in simulated tumours (Supplementary Methods).

#### 2.4.1 Inferring selection strength in TRACERx tumours

We can test this proposition directly by applying each inference pipeline to the cohort of TRACERx tumours (see Figure 8, left panel). We find that more stringent assumptions about cell competition result in higher inferred selection levels, for both LUAD and LUSC tumours. A pipeline trained on Model 2, which implements direct competition, will infer significantly stronger selection across tumours than a pipeline trained on Model 1, which assumes that selection only infers a cell’s ‘inherent’ fitness properties (Model 1 versus Model 2, LUAD: *p* = 2 · 10^−30^; LUSC *p* = 2 · 8^−20^, using 1-sample two-sided T-test; see Supplementary Figure 21). Using a framework based on Model 3, which assumes cells compete for existing space in a fixed-size tumour, results in still-stronger inferred selection (Model 2 versus Model 3, LUAD: *p* = 10^−6^; LUSC, *p* = 2 · 10^−20^).

This implies that our ability to infer selection from real tumours is strongly influenced by model assumptions. In particular, we note that the strongest selection values are inferred using Model 3, we infer slightly higher selection strength from LUSC than LUAD tumours, consistent with previous observations that LUSC tumours have more regional subclonal expansions [21].

We conclude that using Models 1 and 2, in which competition for survival is weaker and cells in existing tissue can never be replaced, introduces a bias towards neutral forms of evolution. These poorly-calibrated, weak-competition models may become a ‘self-fulfilling prophecy’: when models are not carefully calibrated to the underlying properties of lung cancer, the resulting inferences may underestimate the influence of selection on tumour growth.

## 3 Discussion and Conclusion

Over the last fifteen years, extensive genetic intra-tumoural heterogeneity has been detected in a variety of cancers [23, 24]. This in turn has fuelled debate over the role of selection in tumour development, as researchers have attempted to determine whether this ITH reflects fitness differences between cell lineages or develops as a result of neutral diversification [16, 25, 26]. Computational models have become an increasingly popular tool for investigating the rules which govern tumour growth [18, 5, 11, 12], and in particular for distinguishing between the presence and absence of genetic selection [10, 8]. However, recent research has made it clear that different models are necessary to capture the properties of different tumour types and thus make reliable cancer-type-specific inferences [9].

In this work, we re-examined an assumption to common to almost all models in the literature: that fitter cells simply divide more and die less, and compete only indirectly for space and resources. This encodes a very strong limitation about the way that selection ‘works’, which may be relevant in colorectal cancers but may not be appropriate for NSCLCs, where regional subclonal expansions and clonal illusions are common [21]. We investigated whether models which involve ‘active competition’, in which a cell’s proliferation and survival is directly affected by the fitness of its neighbours, are better able to replicate the properties of lung cancers.

In order to replicate the clonal illusions seen in real lung tumours, we find that it is necessary to make much more stringent and computationally expensive assumptions than have been included in any previous tumour-evolution ABM. Our best-performing model (Model 3) assumes not only that cells compete actively within local ‘demes’, but that demes themselves compete for survival when the tumour has already reached full size, with fitter groups of cells able to kill and replace their less-fit neighbours. These mechanics should be included in future models of lung cancer, especially for those used to identify selection pressures in the clinic. They should also be considered when modelling subclonal expansions in other solid cancers (for example, those observed in the breast [27]).

These results also have implications for attempts to identify and predict the lineages which drive metastatic spread [28]. Recent analyses of single-cell NSCLC data have suggested that subclones which have higher potential to seed metastases are also those which *divide* more [29]. Our findings imply that this enhanced proliferation may be linked to invasiveness, in that these fitter cells both divide more *and* kill off their neighbours to spread through existing regions of a tumour. This invasiveness would enhance the ability of highly proliferative clones to colonise normal tissue and seed metastatic lesions. This link may also partly explain why some forms of genomic ITH have been linked to poorer patient prognosis [21]. Further experimental work is necessary to confirm these findings and investigate their full implications.

We also sought to quantify the impact of model design choices on our ability to infer selection strength from real tumours in the TRACERx cohort, using Approximate Bayesian Computation and machine learning. We found that, when using an inference pipeline based on this ‘two-stage, direct competition’ model, we inferred significantly stronger selection across the cohort (in both LUAD and LUSC tumours) than when using a simpler ‘indirect competition’ model (loosely based on SCIMET [11], with non-uniform mitotic age and a saturating effect of selection on proliferation). This suggests that the assumptions of tumour evolution models may indeed be ‘self-fulfilling’: models which assume very little direct competition between cells will tend to underestimate the strength of selection. These results underline the importance of closely matching agent-based models to the properties of the underlying tumour. We also note that this analysis is made possible only by multi-region sequencing data (which allows us to identify clonal illusions). The tumour mutational burden alone is insufficient to distinguish poorly-fitting models (Model 1 and 2) from our best-performing model (Model 3).

As with all modelling-based analyses of tumour growth, our study is limited by computational power and a lack of reliable estimates for some parameters. In particular, our ability to distinguish neutral and high selection is likely dependent on our assumptions about the age of the tumour (1 year) and the size of the local environment (10,000 cells). Whilst these assumptions are clinically reasonable, they are also determined by computational constraints, which prevent us from conducting full sensitivity analyses for all fixed parameters. Generating tens of thousands of simulations of much larger, older or more spatially granular tumours is not currently computationally feasible.

Similarly, the three models we discuss here do not cover all possible sets of assumptions about the mechanisms of selection in tumours. We have focused on what we believe to be the key distinctions between forms of intra-tumoural competition (direct versus indirect competition for resources, competition for new space versus occupied space), and avoided introducing extra parameters where possible. Other distinctions should be examined when modelling different cancers: for instance, models of colorectal cancer growth often incorporate cell differentiation and focus on comparing models of asymmetric and symmetric stem cell division.

Our ‘two-stage, direct competition’ model (Model 3) improves upon the ability of the current literature to replicate the properties of NSCLC, and is (to the best of our knowledge) the first three-dimensional model of lung cancer scalable enough to draw tumour-specific inferences. However, it is only able to produce numbers of clonal illusions on the lower end of the appropriate range. Our models also tend to overestimate the number of mutations shared between regions, suggesting that overall we are still underestimating the full extent of regional diversification in lung tumours. Further work should focus on addressing this discrepancy, perhaps by leveraging greater computational power than was available here to increase tumour age or include competition with normal cells (as in the work of Noble and coworkers [9]). This work demonstrates the mechanistic insights that can be gleaned from tumour-type-specific models of cancer evolution. Future simulation studies should carefully examine their hard-coded assumptions about inter-cellular interactions, and benchmark the properties of their model tumours against real data. This work establishes a precedent for these tumour-type-specific inference models, and should motivate researchers to conduct these benchmarks as a matter of routine. This is a vital first step to the incorporation of computational models into clinical practice.

## 4 Methods

Here we detail the general principles that govern the models presented in this paper. The full details of individual models are provided in the supplement.

### 4.1 Mutation, selection and death

All simulated tumours begin as a single transformed cell. We are interested in modelling only the acquisition and spread of mutations during tumour evolution. As a result, we ignore mutations accumulated prior to tumorigenesis and assume this founding cell is ‘blank’, i.e. contains no mutations.

When a cell divides, each daughter cell acquires a number of exome mutations, *n*_*µ*_ ≥ 0; *n*_*µ*_ is Poisson distributed with mean *µ* = 0.4 (towards the lower end of the range provided in [30]). We model all mutations as SNVs and do not consider copy-number or structural rearrangements. We use the infinite-sites assumption [31], so that no mutation can occur more than once or be lost after it has been acquired, following well-established convention [10, 25, 6, 8].

Somatic mutations are divided into two categories. A mutation is a ‘driver’ with probability *θ* = 10^−5^ (a common literature estimate originating with the work of Bozic *et. al*. [32]), and confers a fitness boost of strength *s*. (We assume that cells can acquire at most one new driver mutation on each division, such that if a cell which has acquired *n*_*µ*_ new mutations, one of them is a driver with probability *n*_*µ*_*θ*.) All other mutations are neutral, on the assumption that positive selection strongly outweighs the deleterious effects of passenger mutations [33].

As cells evolve, they acquire driver mutations, which confer boosts to cell fitness. These boosts have size *s*, a simulation parameter referred to as the selection strength. These fitness boosts always affect proliferation-fitness, but only affect cell *survival* in some models. In all models, proliferation-fitness evolves as follows.

If we assume that *p*_0_ is the probability that the founding cell will divide in a given 1-day timestep, we assume that cell with *k* drivers will divide with a per-timestep probability

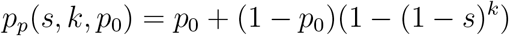

The function is equal to *p*_0_ where a cell has no drivers, or where those boots are ineffectual (i.e. for *s* = 0 or *k* = 0). As the number of boosts increases, however, the probability of division approaches 1 asymptotically. This ensures that each additional boost has a (diminishing) effect on proliferative fitness, whilst preventing nonsensical probabilities of division (i.e. *p* > 1).

This probabilistic division scheme allows cells to have variable mitotic ages– an improvement on existing three-dimensional simulation frameworks designed for tumour-specific inference [11, 12], which assume that cells either divide or die at every timestep and so force all cells alive at the *n*th timestep to have divided exactly *n* times. This will result in an unrealistically uniform accumulation of mutations across the population.

The fitness effects of driver point mutations are generally considered to be in the range 1-10% [34, 11]. In this study we explore selection values in the range 0 ≤ *s* ≤ 0.1. All drivers in a tumour have the same effect on cell fitness.

### 4.2 Deme structure and cell migration

In all simulations we assume that cells are well-mixed within ‘demes’ with a maximum capacity of *N*_0_ = 10, 000 cells [10, 11, 12] for the sake of computational efficiency. Inter-deme competition (represented in models 2 and 3) scales with the number of demes in the model and so is made cheaper by coarser spatial graining (i.e. larger deme sizes). This limit ensures that even in the smallest model considered here (e.g. in the third model presented, which has a final tumour size of 5 million cells), an individual deme represents only 0.2% of the tumour’s population.

Demes are arranged on a three-dimensional grid, governed by a Moore neighbourhood with 26 neighbours. When a deme splits, half of its population moves into a chosen neighbouring deme at random. Cells in demes which are completely surrounded (i.e. which have no empty neighbouring demes) do not divide, as some level of contact with normal tissue is assumed to be necessary for proliferation.

### 4.3 Sampling tumours

At the end of the simulation, we take 8 simulated samples (the maximum number obtained from any tumour in the TRACERx dataset [21]), each comprising 1% of the tumour’s surface cells (i.e. 1% of active demes). We ensure that samples are as distant from each other as possible by randomly generating 20,000 possible combinations of 8 central demes, calculating the sum of the total pairwise distances between them, and choosing the set of demes which are furthest apart according to this metric.

Once 8 spatially-separated demes have been chosen, we generate samples from the surrounding tissue. We treat these demes as the ‘central points’ of the sample, and take the 1% of demes closest to them (roughly 5 demes per sample for the smallest Model-3 tumours, and many more for Models 1 or 2). We check that none of these samples are overlapping (i.e. we have not chosen the same deme twice). If we have, we repeat the selection procedure described above, as many times as necessary, until we have 8 spatially-separated non-overlapping samples. If the population of any sample *N*_*i*_ exceeds *N*_*c*_ = 50000 cells, we include each cell in the sample with probability *N*_*c*_*/N*_*i*_, resulting in a sample of roughly 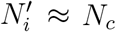 cells. This approximates the size of samples from tumours in the TRACERx cohort [28].

### 4.4 Realistic sequencing of simulated tumours

We assume a tumour purity of 80% and a total read depth of 400x, translating into an ‘effective’ read depth of *R* = 320x.We further assume that our sequencing protocol is unable to detect mutations in a sample if their CCF in that sample is less than 1%. In the TRACERx dataset, real tumours use a threshold which begins at 0.05% and increases as tumour purity decreases. We choose the more stringent end of this threshold largely for computational efficiency, to avoid simulating the sequencing of many low-frequency mutations which will end up being filtered out by the region-specific region thresholds described later in this section.

We assume an error rate of *ϵ* = 0.0001 (0.01%) inherent to the sequencing process. Thus a mutation with identifier *j*, with true CCF *p*_*ij*_ *≥* 0.01 in sample *i*, has a detected CCF

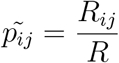

where *R*_*ij*_ is the number of reads of mutation *j* in sample *i*, distributed binomially as *R*_*ij*_ *∼ Bin*(*R, p*_*ij*_(1− *ϵ*)). We further assume that, at this sequencing depth, there is minimal variation in the number of reads at each locus, such that we can assume a uniform *R* = 160 ‘attempts’ to read each mutation. This allows us to avoid further increasing the amount of stochasticity in the model.

We recognise that these assumptions may affect the distribution of observed mutational CCFs in simulated tumours. To minimise the dependence of our model results on the precise shape of this distribution, we choose summary statistics which depend only on whether a mutation is *fixed* (CCF=1.0), *unfixed* (0 < CCF < 1.0) or *absent* (CCF=0.0) in a tumour region.

To calculate summary statistics from *M* simulated samples, we must first ensure these samples are well-separated. The first sample is chosen at random (out of the available 8 samples); the second sample is chosen to maximise its distance from the first; the third sample (if *M* > 2) is chosen to maximise the sum of its distances from the first two, and so on.

Once a set of samples has been chosen, we filter mutations in accordance with TRACERx stringency protocols: mutations which have fewer than 10 reads in all individual samples are excluded (i.e. to pass this step, mutations must have a detected CCF of 6.25% in at least 1 sample).

Finally, before calculating summary statistics, we exclude all truncal mutations, defined as mutations with a detected CCF of 1.0 in all samples. In real tumours, it is impossible to tell whether truncal mutations occurred during tumour evolution or were present in the pre-transformed cell, so we consider only subclonal mutations in order to compare ‘like with like’.

### 4.5 Calculating summary statistics

For a tumour with *M* samples, we calculate the following 2*M* + 1 summary statistics:

1. The number of mutations detected in total, across the tumour.

2. The number of mutations detected in exactly 1, …, *M* samples. (Mutations detected in only 1 sample are often referred to as ‘private’.)

3. The number of mutations only detected in 1 sample, and fixed (i.e. with a detected CCF of 1.0) in that sample.

4. The number of mutations which are fixed in exactly 1, …, *M* − 1 samples. (Mutations fixed in *M* samples are clonal and thus excluded.)

Note that there are no repeating statistics here: a mutation which is fixed in one sample and non-fixed in another (i.e. non-private) will count towards statistic 4 but not statistic 3. We do not include ‘at least’ statistics for this reason: i.e. the number of mutations with an illusion of clonality in *at least* one region (i.e. those fixed in 1 ≤ *m* < *M* samples) can be found by summing all statistics in 4, so would be redundant if included directly.

The fitting criteria described here are summarised in Table 2. A full list of parameters included in each model is included in the supplementary material.

**Table 1:** The mechanistic differences between models presented in this paper.

|  | **Model 1** | **Model 2** | **Model 3** |
| --- | --- | --- | --- |
| Competition within demes | No | Yes | Yes |
| Competition between demes (for new tissue) | Indirect | Direct | Indirect |
| Competition between demes (for occupied tissue) | No | No | Yes |
| Stages of growth | 1 | 1 | 2 |
| Death rate | Variable | Variable | Fixed |
| Final tumour size (in millions of cells) | 500 | 100 | 5 |

**Table 2:** A description of the criteria used in this paper to decide whether each model is a ‘good fit’ to NSCLC TRACERx tumours. Here ‘comparable’ means tumours with the same number of sampled regions.

| Criterion | Type | Model 1 | Model 2 | Model 3 |
| --- | --- | --- | --- | --- |
| Can the model produce the median **TMB** of comparable TRACERx tumours? | Benchmark | Yes | Yes | Yes |
| Can the model produce the median **TMB-priv** of comparable TRACERx tumours? | Benchmark | Yes | Yes | Yes |
| Can the model produce the median number of **clonal illusions** of comparable TRACERx tumours? | Benchmark | No | No | Yes |
| Across summary statistics, which model has the highest fraction of simulations which fall within the 5th and 95th percentiles of summary statistics for comparable TRACERx tumours? | Relative |  |  | x |
| Which model produces the most ‘confident’ predictions for TRACERx tumours? | Relative |  |  | x |

## Supporting Information

Supporting Information is available from the Wiley Online Library or from the author. Sequencing data from the TRACERx study, used to benchmark the models, are available in the Zenodo repository https://zenodo.org/records/7649257. All simulation code is available in the Github repository hcoggan/Age based-simulations-of-lung-tumour-evolution.

## Acknowledgements

H.C. was supported by a grant from the Engineering and Physical Sciences Research Council (EP/W523835/1). C.M.-R. was supported by the Rosetrees (M630) and Wellcome trusts. N.M. receives funding from Cancer Research UK (CRUK) (DRCPFA-Nov23/100003) and has received funding from the Wellcome Trust and the Royal Society (211179/Z/18/Z) relevant to this work. N.M. also receives funding from Cancer Research UK Lung Cancer Centre of Excellence, Rosetrees, and the NIHR BRC at University College London Hospitals. J.F. receives funding from NIHR BRC at University College London Hospitals.

