## Supplementary Materials for "Agent-based simulations of lung tumour evolution suggest that ongoing cell competition drives realistic clonal expansions"

October 2025

The document details computational methods relevant to the work described in this paper, and contains figures supplementary to the main analysis.

#### 1 Model design

Here we provide full details of individual models presented in the main text. The full code for each model is available in the Github hcoggan/lung-modelling-sims.

##### 1.1 Relating cell cycle times to division probability

A cell with proliferative fitness $f_p$ will divide in a timestep $\Delta t$ with probability $p_p = f_p\Delta t$. Throughout the simulation we work in timesteps of $\Delta t = 1$ day.

Our initial cell has a cell cycle time of 24 hours. This is on the lower end of literature estimates for NSCLC cell lines [1; 2], on the assumption that cells are less well-adapted and thus slower to divide in the early stages of tumour evolution. This corresponds to an initial proliferative fitness (probability of division per day) of $p_0 = 0.631$ for the founding cell.

We calculate this as follows. If a cell divides in a timestep $\delta t$ with probability $\alpha\delta t$, then the probability that it has not divided at time $t$ is governed by

$$p(t + \delta t) = (1 - \alpha\delta t)p(t)$$

In order not to have divided at time $t + \delta t$, it must not have divided up to time $t$, and must continue not to divide in the following timestep. As $p(0) = 1$, this gives

$$p(t) = e^{-\alpha t}$$

The probability per unit time, $f(t)$, that a cell will divide between time $t$ and time $t + \delta t$ is given by

$$f(t)\delta t = \alpha\delta t \cdot p(t)$$

and so the probability density describing a cell's likelihood of dividing at time $t$ is simply

$$f(t) = \alpha e^{-\alpha t}$$

The cell cycle time is simply the average time at which a cell divides, and is given by

$$\tau = \int_0^{\infty} t f(t) dt = \frac{1}{\alpha}$$

To calculate the probability that a cell will divide within a timeframe $\Delta t$, $f_0\Delta t$, then, we simply ask what fraction of the cells *have not divided* by time $\Delta t$:

$$f_0\Delta t = 1 - p(\Delta t) = 1 - e^{\Delta t/\tau}$$

A cell cycle time of $\tau$=1 day and a timestep of $\Delta t = 1$ day suggests that we expect roughly $63\%$ of cells to have divided in each timestep, so our baseline fitness becomes $f_0\Delta t = f_0 = p_0 = 0.631$.

#### 1.2 Defining Model 1: The 'indirect competition' model

Our first model assumes that selection affects only a cell's likelihood of division, and that cells compete only indirectly for space and resources. Within demes, cells have no impact on each other's outcomes (division or survival). We use this model to test whether the properties of lung cancer, in particular recent subclonal expansions, can be replicated in a model that assumes fitness is an inherent property of a cell. The model can be thought of as an adaptation of SCIMET [3; 4; 5] with non-uniform mitotic age and saturating selection effects.

##### 1.2.1 Division and death

As in all models, driver mutations increase a cell's probability of division asymptotically towards 1 (as in Equation **??**). Cells die with a fixed baseline probability $\lambda_0$ per timestep; this death rate is a variable simulation parameter. Cells with more driver mutations tend to die less, so that a cell with $k$ driver mutations (with selection strength $s$) dies with a per-timestep probability

$$\lambda(s, k) = \lambda_0(1 - s)^k$$

Cell death does not occur if demes are at 10% capacity or less (i.e. $N \leq 0.1N_0$). As tumour growth is driven by full demes splitting in half (see below in Section 1.2.2), these 'mostly empty' demes are extremely unlikely except in the very early stages of tumour growth, when only one deme is occupied. We impose this rule to avoid the stochastic death of all tumour cells when the tumour is still very small. Mechanistically, this represents saturating competition with the environment: cells in a largely empty deme have enough resources to divide without dying until the population breaches some threshold (here 1000 cells). Division and death are both probabilistic, which ensures variable mitotic age: in any given timestep a cell need neither divide nor die.

##### 1.2.2 Cell migration and tumour growth

When a deme becomes full (i.e. $N > N_0$ after division and death), its population splits in half and colonises a randomly-chosen empty neighbouring lattice-point, if one exists. This creates an implicit inter-deme competition for space, with fitter cells dividing faster and colonising space that less-fit cells might otherwise have used. Less fit cells are likelier to be trapped in the centre of the tumour and are less likely to be sampled at the end of the simulation. At every timestep, demes are iterated over in a random order, with cells dividing and dying probabilistically and demes splitting where they are over capacity. This has the effect of randomly breaking ties between equally-fit demes competing for the same empty neighbouring lattice-point at the same timestep.

##### 1.2.3 Computational constraints

The relatively simple assumptions about competition in this model mean that it is able to achieve a final tumour size of roughly 500 million cells. The time taken to reach this size varies, depending on the baseline death rate $\lambda_0$. The average fractional growth rate $\epsilon$ of the founding lineage is given by $1+\epsilon = (1+p_0)(1-\lambda_0)$. As division and death are both stochastic, setting $\epsilon$ too close to zero results in tumours that stall or shrink, and never reach full size; experience suggests that a minimum growth rate of around $\epsilon = 0.01$ is necessary to avoid this. This corresponds to $\lambda_0 = 0.38$, which we use as our maximum death rate. We explore death rates $\lambda_0 = [0.0, 0.1, 0.2, 0.3, 0.35, 0.38]$, which will result in tumour ages of between forty days and five years. (These calculations are based on the properties of the founding cell, and will necessarily be overestimates if driver mutations sweep the tumour and increase the overall growth rate; we take them as a guideline here.)

#### 1.3 Defining Model 2: The 'direct competition, continual-expansion' model

In the second model, we assume that cells compete both within and between demes. Competition between cells is for survival, so that a cell's probability of dying is no longer dependent only on its own genotype but on its surrounding ecology. Demes compete based on their relative fitness to move into empty space, instead of expanding only when they are full. The tumour continually expands, and tissue that has been colonised can never be *recolonised*. We use this model to test whether the properties of lung cancer can be replicated by a model that assumes that a cell's fitness is relative within a continually expanding tumour, i.e. that there is no competition for existing space.

##### 1.3.1 Division and death

A cell's proliferative fitness (i.e. its probability of dividing per timestep) depends on its driver mutations exactly as in 1.2.1.

Here a cell also has a 'survival-fitness', given by $f_s = (1+s)^k$. This increases without bound with selection strength $s$ and number of driver mutations $k$, as it is only ever measured relative to the fitness of other cells. When a deme is at more than 10% capacity, cells within it compete for survival: this encodes the idea that, in an environment with limited resources, cells are less likely to survive if there are a large number of fit cells in their vicinity. For a baseline death rate of $\lambda_0$, a cell's probability of dying in a given timestep is given by

$$\lambda = \max(1 - (1 - \lambda_0)\frac{f_s}{\bar{f_s}}, 0)$$

where $\bar{f_s}$ represents the average fitness of all cells in the deme. Cells die with probability $\lambda_0$ if their fitness is equal to the average fitness of cells in their local environment; a fitter cell $(f_s > \bar{f_s})$ is more likely to survive. This eliminates the possibility of true 'asymptotic selection', as survival probabilities are now interdependent. In Model 1, as cells gain driver mutations, their probabilities of both survival and proliferation increase asymptotically towards 1 at every timestep, and there will come a point at which further driver events 'stop mattering': if all cells in a deme have some large number $k$ of drivers, a cell with $k + 1$ drivers is no longer meaningfully advantaged. In Model 2, survival probabilities are calculated from *relative* fitnesses, so the emergence of a cell with an extra driver will force up the death rates of all other cells and decrease its own, regardless of the baseline number of drivers $k$. This avoids a scenario in which selection pressures eventually 'die off', and evolution becomes neutral: in Model 2, cells can always get fitter.

Cells divide and die until their demes are over-capacity $(N > N_0)$ at the end of a timestep, at which point both processes stop until the next 'splitting event', described below.

##### 1.3.2 Cell migration and tumour growth

If cells compete for survival, the population of a deme is no longer a reasonable proxy for its average population fitness. In Model 1, fitter cells divide more and die less, so if one deme reaches capacity before another, we can reasonably say that the cells within that deme have 'outcompeted' their neighbours. A 'first-come, first-served' approach to deme splitting, with over-capacity demes immediately claiming nearby empty space, therefore represents implicit competition for space and resources that fitter cells would be expected to win.

In Model 2, competition for survival decouples the relationship between the overall death rate in a deme and the fitness of its cells. As fit lineages emerge, they increase the likelihood that their less-fit neighbours will die, which might cause the population of the deme to drop even as its average fitness rapidly increases. As a result, we cannot allow demes to split when they become full if we want 'fitness' within the simulation to have any meaning at all (i.e. if we maintain that increasing fitness should increase the likelihood that a cell's descendants will be sampled from the tumour surface). We therefore assume that demes split *all at once*, when the tumour as whole reaches a threshold cell density and runs out of space to contain its growing population. (We could alternatively assume that demes split individually when they become a certain amount fitter than their immediate neighbours, but this would prevent growth in neutral tumours.)

We implement these splitting events when all active demes (those with empty neighbours) are over-capacity. The 'splitting algorithm' works as follows:

1. Identify all empty demes which neighbour *at least one* full deme.
2. Iterate over those empty demes in a random order. For each empty deme:
   (a) Identify all full demes neighbouring this deme. If there are none remaining, move on to the next empty deme.
   (b) If there are full neighbouring demes, choose one to split into the empty deme. The probability of choosing deme $i$, where the cells have average survival-fitness $\bar{f}_s^i$, is $\frac{\bar{f}_s^i}{\sum_j \bar{f}_s^j}$, where the sum over $j$ in the denominator encompasses all demes competing for the empty deme. This is to say that demes compete for space and resources. Each deme has a chance of success linearly proportional to its relative fitness, reflecting the dynamics of competition between cells. Of course, if there is only one full deme trying to expand, then this will be chosen regardless of its fitness.
   (c) Move each cell from the full deme into the empty deme with probability ½.
3. If a deme is still full at the end of this simulation, mark it as 'inactive' and stop tracking cells within it. All empty demes identified in step 1 are either no longer empty or have no remaining full neighbouring demes. Thus, any deme still *full* must have been outcompeted and cannot have any empty neighbours remaining, and so has no chance of splitting at a future timestep. The cells within it will not be sequenced at the end of the simulation, when samples are taken from the tumour's surface.

After the splitting event, division and death resumes. This continues until the tumour reaches full size.

##### 1.3.3 Computational constraints

Competition both between cells and between demes requires repeated comparisons at every timestep, which increases the computational demands of this model. As a result, we are limited to a final tumour size of 100 million cells. We explore death rates $\lambda_0 = [0.0, 0.1, 0.2, 0.3, 0.35, 0.38]$ as in Model 1.

#### 1.4 Defining Model 3: The 'two-stage, direct competition' model

In our final model we assume that cells compete both within demes (for survival) and between demes (for both empty and occupied space). Tumour evolution proceeds in two stages: one 'exponential' and one 'homeostatic'. This is an idealisation of the general principle that growth slows as the tumour size increases, and that cells in a large tumour compete more fiercely for space and resources.

In the first 'exponential' stage, cells divide without dying, and demes split when full. At the first timestep at which the population of the tumour exceeds 5 million cells, tumour growth stops. We assume during the second 'homeostatic' stage that the tumour has run out of space to grow into, and that an increase in local cell density triggers competition, not expansion. Cells spread through the tumour by out-competing their less-fit neighbours for space and nutrients; cells which lose such competitions die, and yield their resources to fitter, invading lineages.

We use this model to test whether allowing cells to invade occupied demes and replace less fit lineages is sufficient to reproduce the properties of lung cancer. This two-stage approach is adapted from the work of Zhao and coworkers [6], who assumed that cells in colorectal cancer glands continue to age (i.e. divide and die) after the maximum number of glands has been reached, but has not been deployed in any three-dimensional model of cancer growth to our knowledge.

##### 1.4.1 Division and death

In all cases, cells proliferate as described in Section 1.2.1. During the first stage of growth, there is no cell death. During the second, demes which are over capacity $(N > N_0)$ after the division step are forced to compete for survival, as it is assumed they have run out of space. As in Model 2, a cell with $k$ drivers of strength $s$ has survival-fitness $f_s = (1+s)^k$. A cell with survival-fitness $f_s$ survives with probability $p = \min(1, \frac{N_0 f_s}{F})$, where $F$ is the total fitness of all cells in the deme, and dies otherwise. If the deme contains $N$ cells of equal fitness, then their probability of survival will each be $N_0/N$, and we will expect roughly $N_0$ cells to survive at the end of the step. Thus, cells in a deme compete on fitness for sufficient space and resources. Over time, fitter lineages should slowly take over the population of a deme. This assumption means the simulation has no independent death rate parameter $\lambda_0$.

##### 1.4.2 Cell migration and tumour growth

During the first stage of tumour growth, demes split into empty neighbouring demes when full, as there is no competition and so fitter cells merely proliferate faster. After a split, both demes are then under-capacity and proliferation can resume. This expansion continues until the tumour's total population exceeds 5 million cells. (This stage of tumour growth can be thought of as a smaller version of Model 1 with no cell death, such that $\lambda_0 = 0$.)

In the second stage, in order to allow fitter lineages to spread beyond the demes they arise in, we also implement competition *between* demes. At the end of each timestep, the populations of fitter demes split in half (i.e. each cell moves with probability $0.5$ and stays otherwise) and replace the populations of their less-fit neighbours. We assume that all cells in the invaded deme die and disintegrate immediately, so that space and nutrients are available to the cells replacing them.

The process of replacement is probabilistic. If deme $i$ is a neighbour of deme $j$, and the average survival-fitnesses of their populations (before the replacement step) are $\bar{f}_s^i$ and $\bar{f}_s^j$ respectively, then the probability that deme $j$ will be become available to be replaced by deme $i$ is

$$p(i \to j) = \begin{cases} 1 - \frac{\bar{f}_s^j}{\bar{f}_s^i} & \text{if } \bar{f}_s^i \geq \bar{f}_s^j \\ 0 & \text{if } \bar{f}_s^i < \bar{f}_s^j \end{cases}$$

This is a parameter-free function designed to ensure that less-fit demes will never replace fitter demes, and that replacement-availability becomes certain $(p(i \to j) \to 1)$ as $\frac{\bar{f}_s^j}{\bar{f}_s^i} \to 0$, i.e. as deme $i$ becomes infinitely fitter than deme $j$. If deme $i$ has several demes available to be replaced by it in a single timestep, then it chooses one of these neighbours at random. Each deme can only be replaced, or replace another deme, once in a timestep, though a deme that has invaded another may itself be invaded by a fitter deme later in the same timestep. As in the expansion stage, demes are iterated over in a random order, so that ties between two demes competing to expand into the same less-fit neighbouring deme are broken at random.

A deme which splits randomly in half to replace a neighbouring deme should not see any changes in its average fitness; a deme which is replaced should see its fitness increase. This means that the process of replacement is inherently self-limiting. Once a deme has replaced its neighbour, the two populations should have similar fitnesses, and so neither will be likely to replace the other until a fitness differential has built up between them over many timesteps of separate evolution.

As before, proliferation only occurs in demes with at least one empty neighbouring lattice-point. The second stage of this simulation therefore takes place entirely on the tumour surface. Surrounded demes are assumed to be low in nutrients and high in mechanical pressure, so living cells do not compete to invade them. Lineages arising during this stage of evolution can colonise the entire tumour by spreading through surface demes.

##### 1.4.3 Computational constraints

The process of replacement means that tumours continue to evolve whilst being kept at a fixed size. This breaks the traditional link between size and age in computational models of tumour evolution [4; 5; 6; 7; 8], and allows us to simulate the dynamics of cell competition in a fixed-size tumour. However, this stage of evolution is very expensive, and scales more than linearly with the number of cells in this tumour. To keep our model computationally feasible, we set the fixed size to be 5 million cells.

The model proceeds for 365 timesteps, or one simulated year. By this point, roughly 1 billion cells should have been produced (assuming 5 million cells undergoing 0.631 divisions per day for around 300 days), though only a fraction survive.

### 2 Inferring selection strength from real and simulated tumours

We use a variant on Approximate Bayesian Computation ([9]) to infer selection strength. ABC has been used by Graham and coworkers to distinguish the presence of zero, one or two selected subclones from single tumour regions [7] and multi-region samples of colorectal cancer [10]; the same team of researchers have used ABC to retrieve selection levels and time of subclone emergence from simulated multi-region tumour data [11]. Curtis and colleagues have also used ABC to deduce differentiation patterns from primary CRCs [6; 8; 12] and seeding times from paired primary and metastatic samples [4]. If fitting models to a relatively small number of tumours, researchers can use 'sequential Monte-Carlo' ABC (ABC-SMC), in which simulations are continually generated from a decreasing parameter space until they converge on the properties of a real 'target' tumour. Depending on the complexity of the simulation and the identifiability of its parameters, one can also select between models using this method, and fit as many parameters as desired (though each new parameter will increase the dimensionality of the problem, so this will depend on computational constraints).

However, ABC-SMC is necessarily iterative and difficult to parallelise (since we must evaluate the properties of some simulated tumours before deciding which parameters to use to generate the next), and its cost scales with the number of target tumours. Here we have more than 300 evaluable TRACERx tumours, and three complex and highly stochastic simulation frameworks, which limits the dimensionality of our inference problem (ideally we would infer 1-3 parameters). We take the approach of Hu *et. al.* [4] and generate a large set of simulated tumours (tens of thousands) from a uniform prior distribution. For each target tumour, we identify the set of these simulations which most closely represent that tumour's properties, and use their parameters to produce a posterior distribution. In Section **??**, we seek to infer only the selection strength $s$.

In this work we combine both approaches, using a method developed by Jiang and coworkers [13] (Figure 1). We split the cohort of simulations in half, into a training set and testing set. We use the training set train a classifier (here XGB-Regressor) to predict $s$ from a the summary statistics $S_i$ of simulated tumours. We apply this classifier to the real tumour to predict its selection strength, $s'_{pred}(S'_i)$. We then use our test set to construct a probability distribution for the selection strength. The distance of a simulated tumour with summary statistics $S_i$ from the real tumour is given by $D(S_i, S'_i) = (s_{pred}(S_i) - s'_{pred}(S'_i))^2$, i.e. the L2 distance between the *classifier's assess-*

*ments* of the selection strength in the real and simulated tumours. We then choose the 1% of simulated scenarios which are closest to the real tumour. Our strategy amounts to using ABC with an ML-driven distance function, and thus constructing our probability distribution from the properties of the 1% of simulated tumours which a classifier 'thinks' are closest to the real tumour. This allows all the flexibility of ML whilst retaining the benefits of ABC (uncertainty estimation and constrained outputs). Our inference process is described in Figure 1).

### 3 Benchmarking inference on simulated tumours

Before applying this inference process to real tumours, we first benchmark it on simulated tumours. This allows us to assess how well we can infer selection strength 'in theory', because in simulated tumours, the underlying selection strength is known. Each benchmark is carried out on tumours from the appropriate model: we attempt to infer selection in Model-1 tumours using a classifier trained on Model-1 tumours, and so on. In this way we can compare how accurately selection can be inferred from each model, *assuming that the model assumptions are appropriate to the tumour*.

To keep this comparison fair, we discard simulations from Model 1 and Model 2 with death rates $\lambda_0 < 0.38$; we have seen above that simulations with low cell turnover are unable to replicate the mutational burden of lung cancer. This means that we have roughly 20,000 simulations per model, of which 10,000 are used to train the classifier; the choice of the 1% most appropriate simulations thus amounts to choosing the 100 'best' model tumours to replicate each 'target' tumour. We assess each model using leave-one-out analysis: we exclude each simulated tumour in turn from the test set and treat it as the target tumour, attempting to infer its underlying selection using the remainder of the test cohort. Because we know the *true* selection value used to generate it, this allows us to assess the accuracy of the pipeline.

We compare the performance of this inference pipeline across models (top to bottom) in Figure 2. We can assess performance by looking both at the average MSE per prediction (i.e. the mean squared difference between true and inferred selection), and the correlation between true and inferred selection (Pearson's R). Perfect inference will result in an MSE of zero and an R of one.

We find that the inference pipeline performs similarly across all models on simulated data. MSE is lowest (and R highest) in Model 3, which best replicates the properties of TRACERx tumours; this is closely followed by Model 1, in which fitter cells divide more and die less without directly competing. Model 2 is the most difficult framework under which to infer selection, as the effects of selection are saturating; the properties of a Model-2 tumour with $s = 0.06$ are not meaningfully different from those with $s = 0.1$.

Under all models, neutral selection remains difficult to correctly identify: generally most models at $s < 0.01$ are assessed at $s \approx 0.01$. Selection is generally overestimated in the range $0.02 \leq s \leq 0.07$, and underestimated thereafter. Within Model 1 and Model 3, we see a clear peak in TMB at $s = 0.02$; inferred

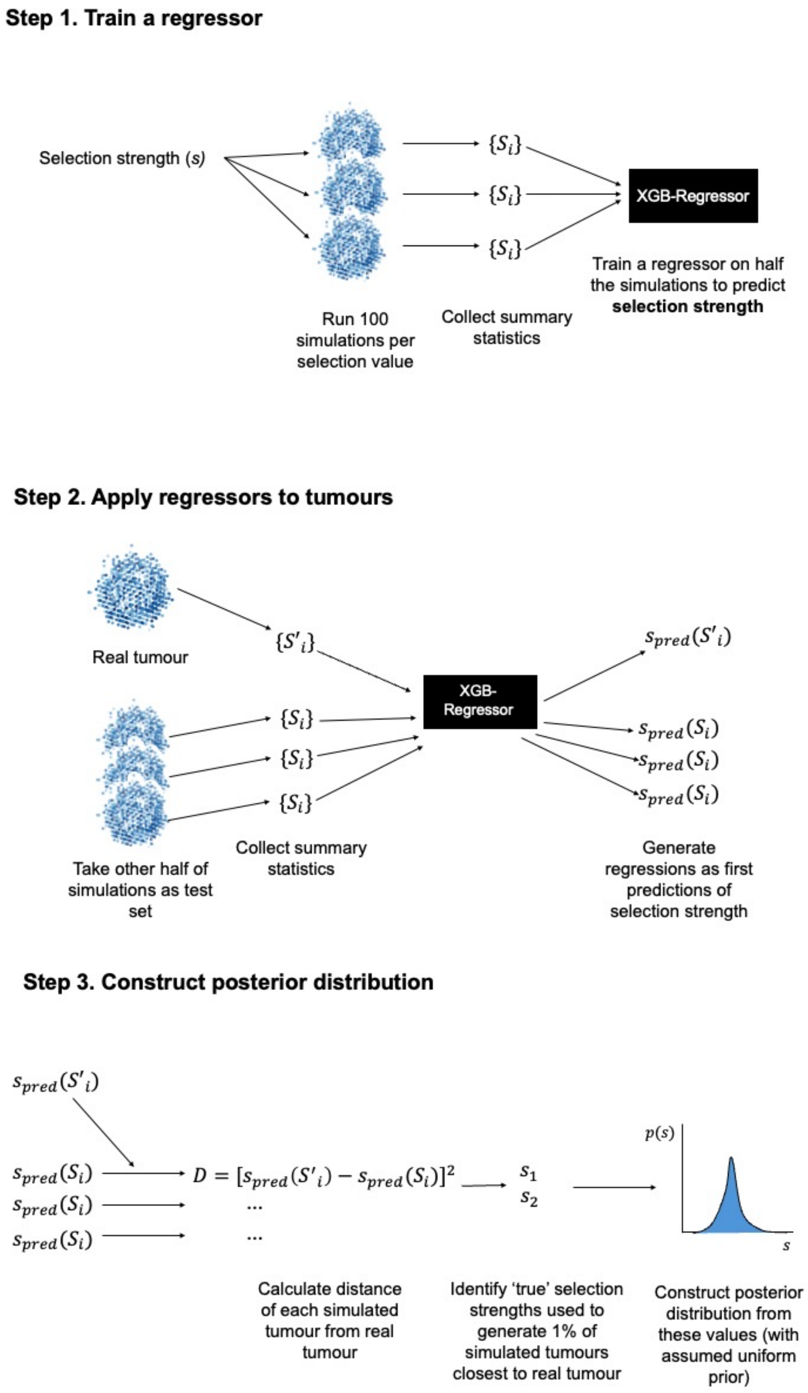


Fig. 1: A schematic of the inference process used to fit continuous selection strength to a real tumour from the TRACERx cohort. Note that the fitting process itself does not involve direct comparison of models, but instead aims to infer the selection strength that would result *if one had already chosen a particular model*.

selection increases sharply around this point, suggesting a reliance on TMB to deduce $s$. We examine more directly later (Figure 3). All models show saturating inference: small differences become increasingly difficult to discern at high selection levels, such that tumours at $s = 0.1$ are usually inferred to have selection levels of $s = 0.07 - 0.08$. For all models, inference becomes increasingly accurate with more samples (see Appendix Figure 20).

This benchmarking process provides a 'best-case scenario' for the quality of inference from real tumours, assuming the inference pipeline is trained on simulations which accurately capture the properties of its target data (here, *in silico* tumours from the same model). As we have seen, these model tumours vary in their resemblance to real lung cancers, and so our ability to deduce selection strength from real tumours will be limited by how well we have modelled the underlying biology. In particular, our choice of model may influence what the classifier 'thinks of' as the properties of high-selection tumours.

Whilst selection is more or less equally easy to infer from all models, the 'decision-making process' learned by the pipeline varies significantly between models. This can be seen by examining the relative importance of each summary statistic to each regressor (as measured by the normalised gain-type importance; see Figure 19). For example, the decision-making process of regressors trained on Model 1 is strongly dependent on the number of samples, $M$. Selection is assigned to low-sample tumours largely based on the overall TMB (Figure 3); when more samples are present, the regressor begins to prioritise the number of mutations fixed in one sample (see Appendix, Figure 19). Regressors trained on Model 2 always make decisions based on the overall number of mutations detected, whereas those based on Model 3 focus on mutations fixed in multiple samples. These differences suggest that the choice of underlying model strongly influences the inference pipeline's idea of what a neutral or high-selection tumour 'looks like', and this in turn may bias our results when selection strength is inferred from a real tumour.

### 4 Pre-processing of TRACERx lung tumour data

TRACERx data is taken from the work by Frankell *et. al.* [14]. To ensure that variations in mutational frequency reflect *inter-tumour* heterogeneity, the dataset excludes samples which initially appeared to have been taken from the same tumour but were later discovered to comprise genomically distinct tumours in the same patient. Tumours have a minimum of 2 and a maximum of 8 samples (see Figure 4). The dataset comprises 382 tumours with a median of 3 sampled regions (inter-quartile range 2.0-4.0).

Mutation CCF values were computed for all mutations detected in TRACERx patients. Mutations were processed as in the TRACERx421 study [14]. CONIPHER [15] was run to compute the cancer cell fraction (CCF) of mutations and to construct mutational clusters. In cases where mutations were removed by CONIPHER during clustering, the mutational CCF in each region was computed by the following formula:

```
compute_mut_ccf <- function(vaf, purity, tum_cn, normal_cn)
```

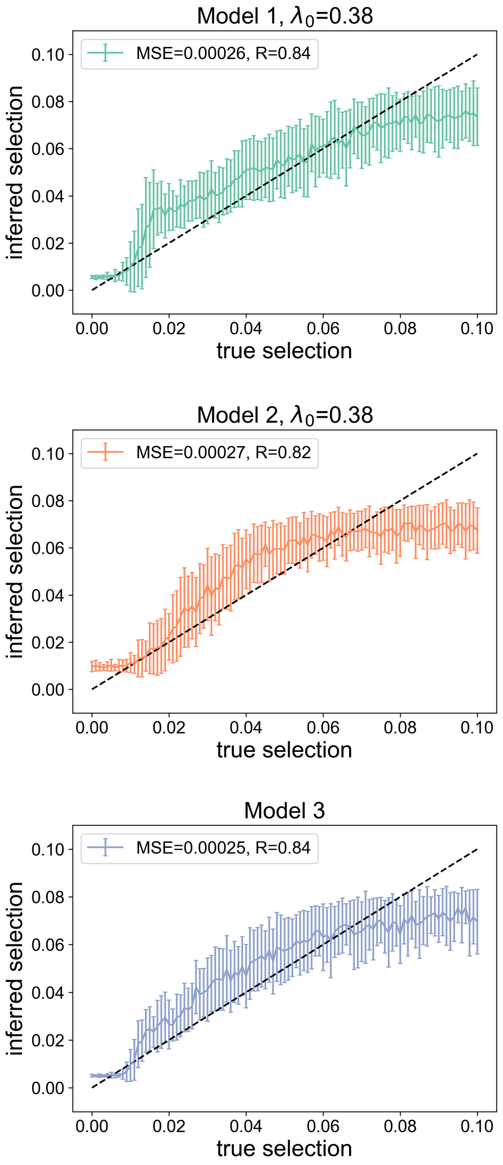


Fig. 2: The relationship between real and inferred selection for simulated tumours in Model 1 (top), Model 2 (centre) and Model 3 (bottom), assuming $M = 2$ samples. Models 1 and 2 have an assumed death rate of $\lambda_0 = 0.38$. The inferred selection is taken as the average of the selection strengths corresponding to the 1% closest tested simulations; error bars indicate the standard deviations of these inferences. The legend indicates MSE (average mean squared error across all predictions) and R (Pearson's correlation between true and inferred selection). The line y=x (corresponding to perfect inference) is plotted in black for comparison. The objective of XGB-Regressor is to minimise the MSE.

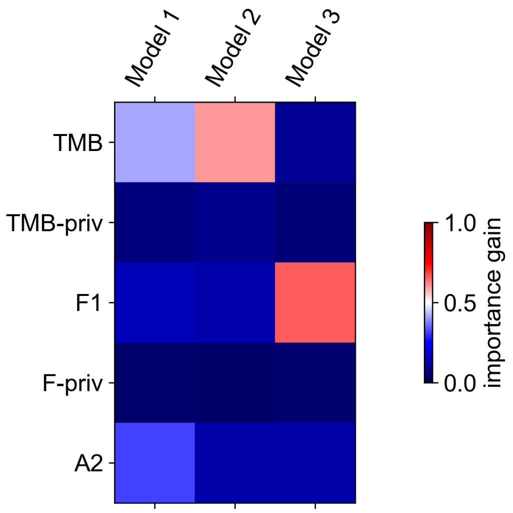


Fig. 3: The relative importance gain for classifiers trained on each of three models, assuming $M = 2$ samples. The $(i, j)$th element represents the relative importance gain of summary statistic $i$ to the regressor trained on *in silico* tumours from Model $j$. Models 1 and 2 have an assumed death rate of $\lambda_0 = 0.38$. Importance gains are normalised across all summary statistics.

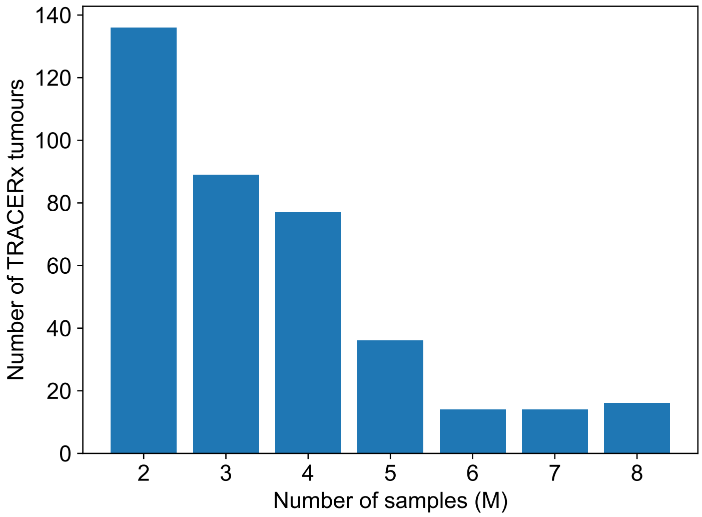


Fig. 4: A bar chart describing the number of samples taken from tumours in the TRACERx cohort.

```
{ vaf/purity * (purity * tum_cn + (1 - purity) * normal_cn) }
```

If a mutation is identified as ‘clonal’ in a particular region by the clustering algorithm used to deconvolve CCF from copy number (CONIPHER, see [15]), we round its CCF to 1.0; if a mutation has a recorded CCF greater than 1.0, we round this down to 1.0. For the purposes of comparison with simulations, we exclude all mutations which are fixed (CCF=1.0) in all regions.

When evaluating model performance (see below), we compare simulated tumours to real NSCLC tumours in the TRACERx cohort. Here we use only

| Parameter | Model 1 | Model 2 | Model 3 | Source |
|---|---|---|---|---|
| Size of tumour | 500 million cells | 100 million cells | 5 million cells | Computational constraints |
| Age of tumour | 6 months-5 years | 6 months-5 years | 1 year | Consultation with colleagues |
| Baseline probability that a cell will die per timestep $(\lambda_0)$ | 0.0, 0.1, 0.2, 0.3, 0.35, 0.38 | 0.0, 0.1, 0.2, 0.3, 0.35, 0.38 | None until deme full | NA (variable parameter) |
| Selection strength | Varied between 0-0.1 | | | [3; 4] |
| Mutation rate (average exome mutations per division) | 0.4 | | | [16] |
| Size of local environment ('deme') | 10,000 cells | | | [4; 16] |
| Size of Moore neighbourhood | 26 cells | | | [17] |
| Size of timestep | 1 day | | | Consultation with colleagues |
| Probability that a mutation is a driver (assuming genetic selection) | 0.00001 | | | [18] |

Table 1: A list of parameters used in each model.

primary tumours with 2 or more samples, taken from patients in which only a single primary tumour was found (i.e. excluding cases in which multiple genomically distinct primary tumours were identified in the same patient). We include all mutations which are trusted by the pass filter (approximately those with 4 reads in the sample of interest and 10 in at least one sample). We exclude mutations which marked as occurring 'early' in tumour evolution (i.e. occurred truncally and before whole-genome doubling WGD, an event that is known to happen early in the evolution of lung tumours [14]).

| Parameter | Model 1 | Model 2 | Model 3 | Source |
|---|---|---|---|---|
| Probability that a mutation is a driver (assuming genetic selection) | | 0.00001 | | [18] |
| Probability that the founder cell will divide, per timestep | | 0.631 | | [1; 2] (see text) |
| Maximum number of samples from any tumour | | 8 | | [14; 16] |
| Minimum CCF required for detection of a mutation in a sample | | 0.01 | | [14] |
| Minimum number of reads required for detection of a mutation in specific sample | | 4 | | [14] |
| Minimum number of reads in any sample required for detection of a mutation | | 10 | | [14] |
| Sequencing depth | | 320x (400x at 80% purity) | | Average read depth [14]; purity set following consultation with colleagues(see text) |
| Sequencing error rate | | 0.0001 | | [16] |
| Cells per bulk tumour sample | | 50,000 | | [16] |

Table 2: A list of parameters used in each model (continued from previous page).

### 5 Supplementary Figures and Tables

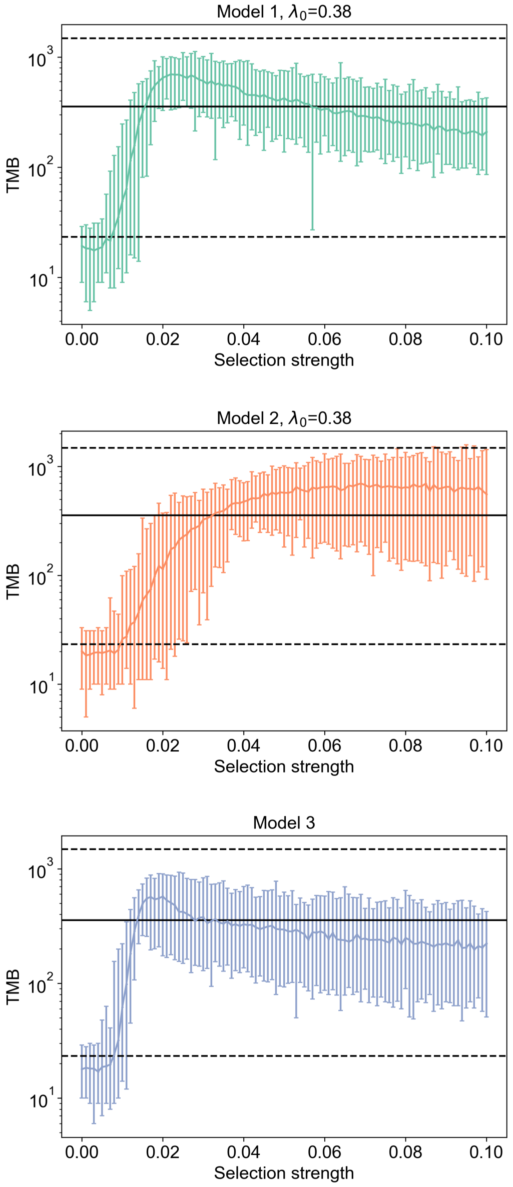


Fig. 5: The variation of simulated TMB in each model with selection strength $s$. Models 1 and 2 have an assumed death rate of $\lambda_0 = 0.38$. We assume $M = 5$ samples. Averages are taken from 100 repeats at each $(s, \lambda_0)$ pair, with maxima and minima indicated by error bars. Dotted black lines indicate the outer limits (5th and 95th percentiles) of corresponding summary statistics from all TRACERx tumours which were sampled at $M$ regions. Solid black lines indicate the medians of these real TRACERx tumour cohorts.

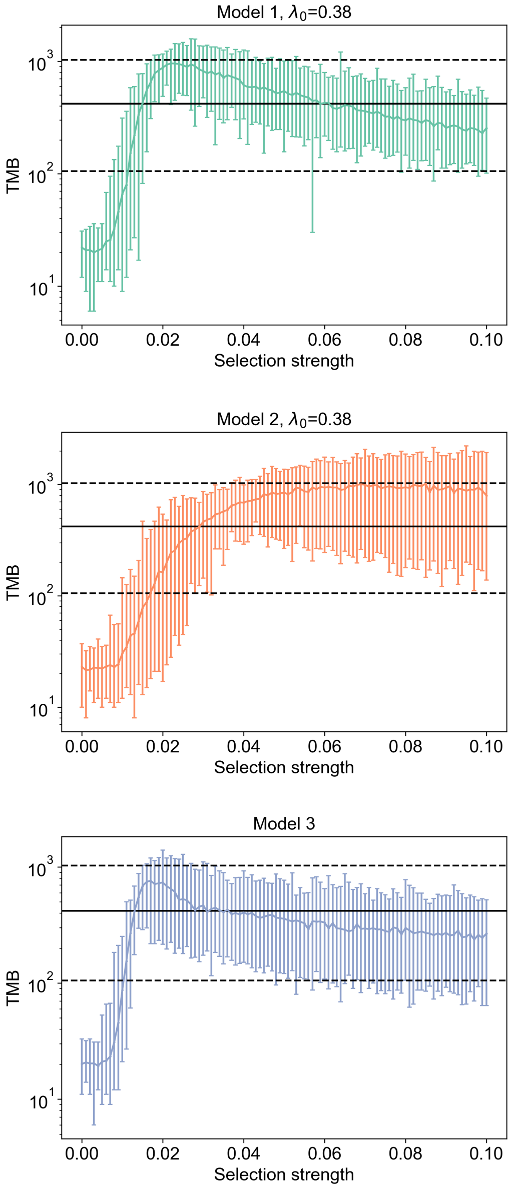


Fig. 6: The variation of simulated TMB in each model with selection strength $s$. Models 1 and 2 have an assumed death rate of $\lambda_0 = 0.38$. We assume $M = 8$ samples.

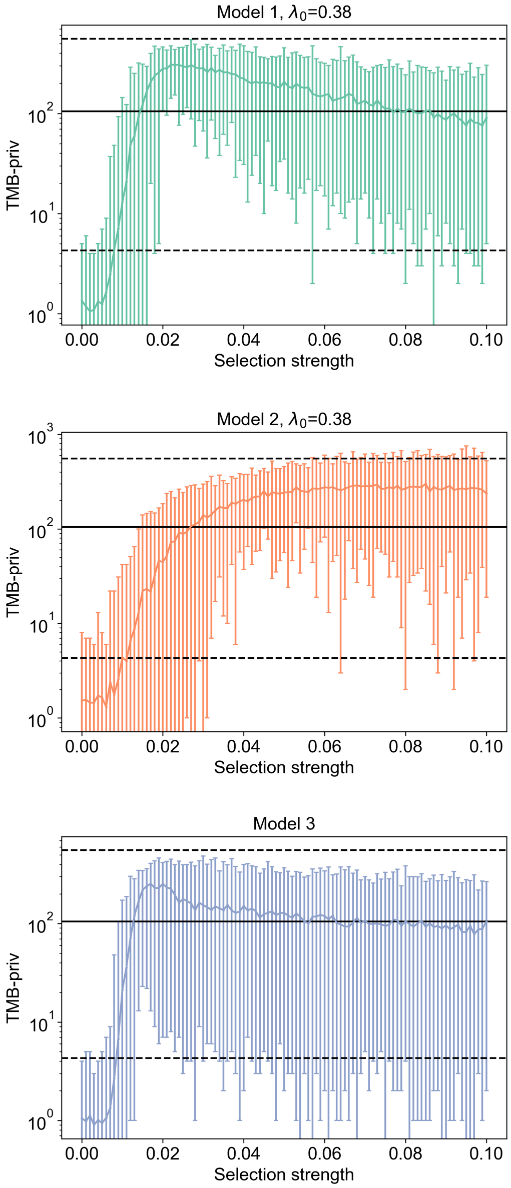


Fig. 7: The variation of simulated *private* TMB (number of mutations only detected in one region) with selection strength $s$. Models 1 and 2 have an assumed death rate of $\lambda_0 = 0.38$. We assume $M = 2$ samples.

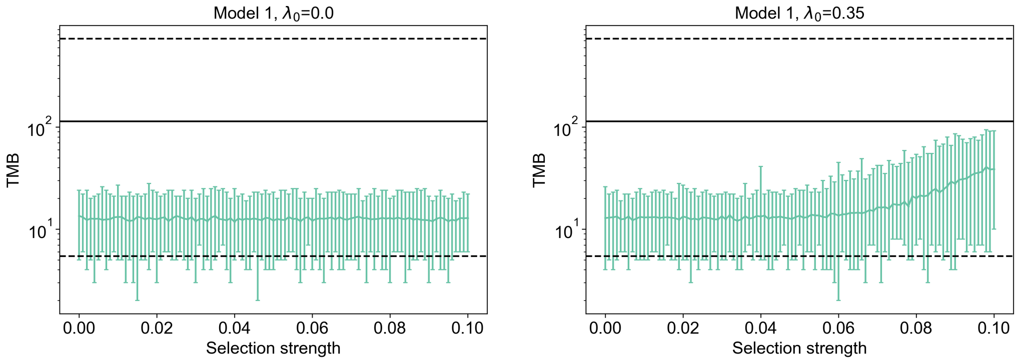


Fig. 8: The variation of simulated TMB in each model with selection strength $s$ in Model 1, assuming a death rate of $\lambda_0 = 0.0$ (left) or $\lambda_0 = 0.35$ (right). We assume $M = 2$ samples.

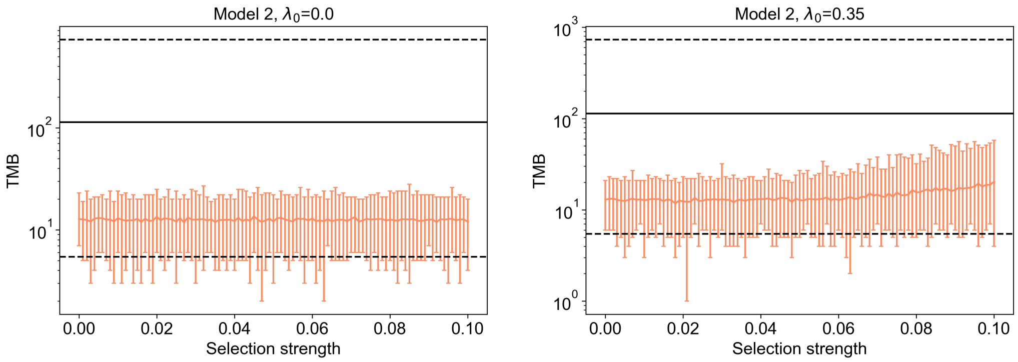


Fig. 9: The variation of simulated TMB in each model with selection strength $s$ in Model 2, assuming a death rate of $\lambda_0 = 0.0$ (left) or $\lambda_0 = 0.35$ (right). We assume $M = 2$ samples.

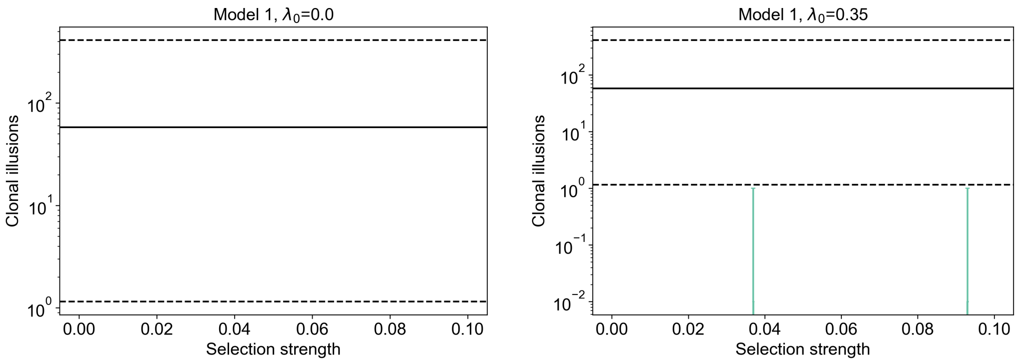


Fig. 10: The variation of simulated CIs in each model with selection strength $s$ in Model 1, assuming a death rate of $\lambda_0 = 0.0$ (left) or $\lambda_0 = 0.35$ (right). We assume $M = 2$ samples.

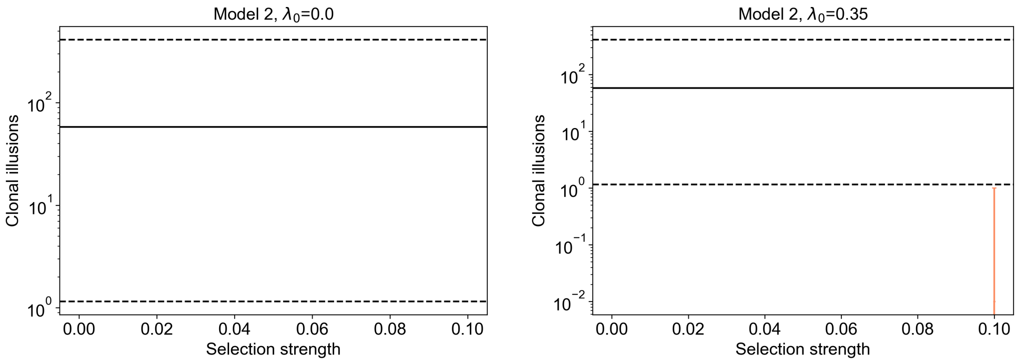


Fig. 11: The variation of simulated CIs in each model with selection strength $s$ in Model 2, assuming a death rate of $\lambda_0 = 0.0$ (left) or $\lambda_0 = 0.35$ (right). We assume $M = 2$ samples.

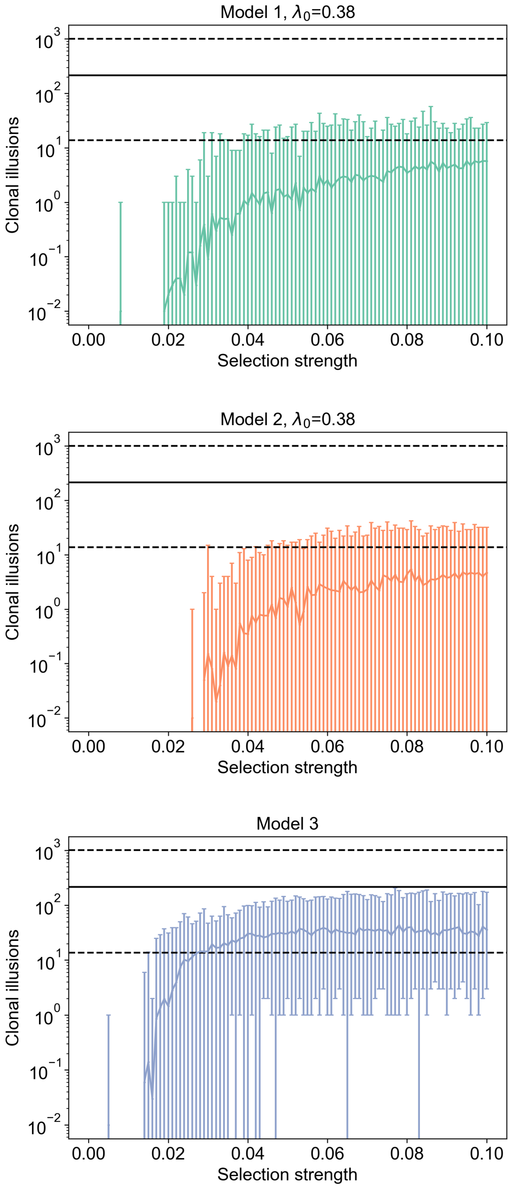


Fig. 12: The variation of simulated clonal illusions (CIs) in each model with selection strength $s$. Models 1 and 2 have an assumed death rate of $\lambda_0 = 0.38$. We assume $M = 5$ samples.

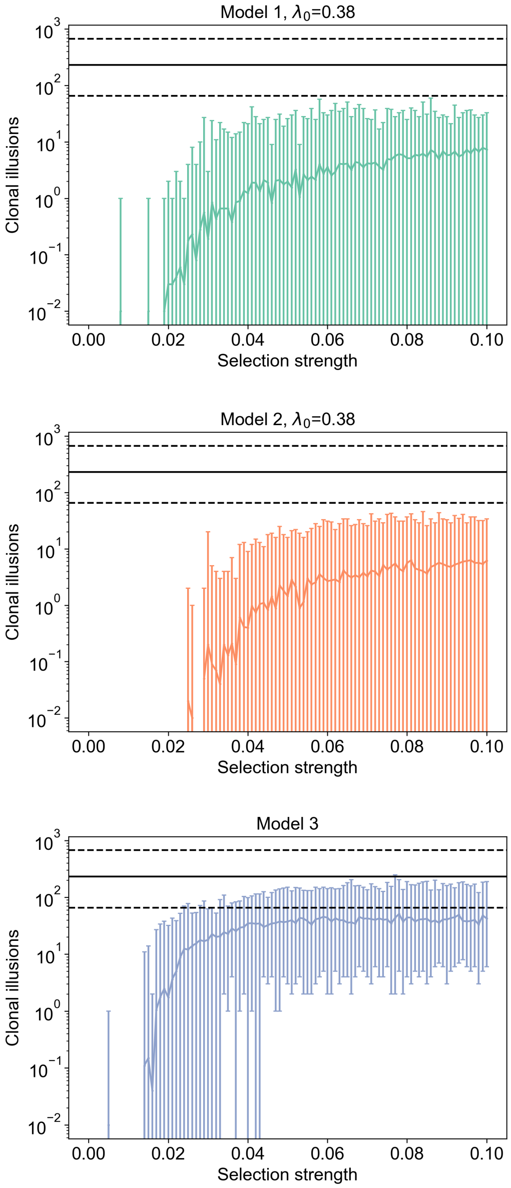


Fig. 13: The variation of simulated clonal illusions (CIs) in each model with selection strength $s$. Models 1 and 2 have an assumed death rate of $\lambda_0 = 0.38$. We assume $M = 8$ samples.

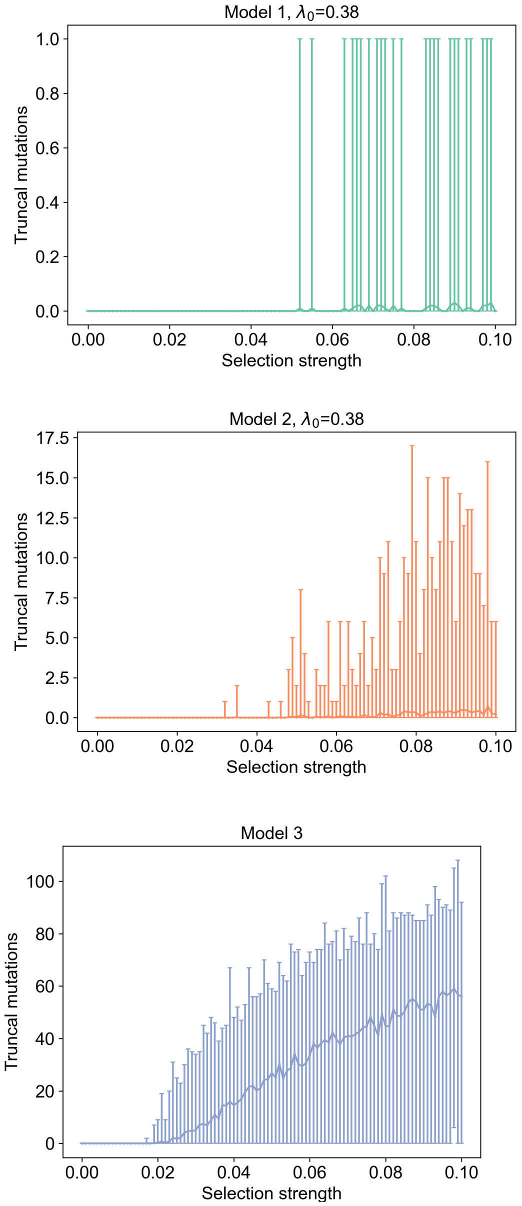


Fig. 14: The variation of the number of truncal mutations in each model with selection strength $s$. Models 1 and 2 have an assumed death rate of $\lambda_0 = 0.38$. We assume $M = 5$ samples.

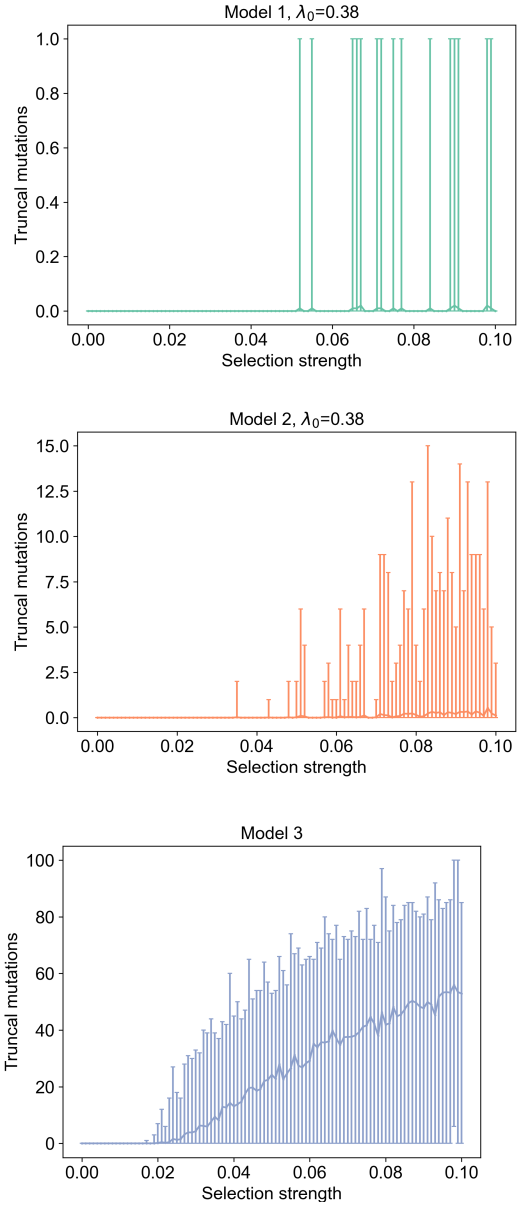


Fig. 15: The variation of the number of truncal mutations in each model with selection strength $s$. Models 1 and 2 have an assumed death rate of $\lambda_0 = 0.38$. We assume $M = 8$ samples.

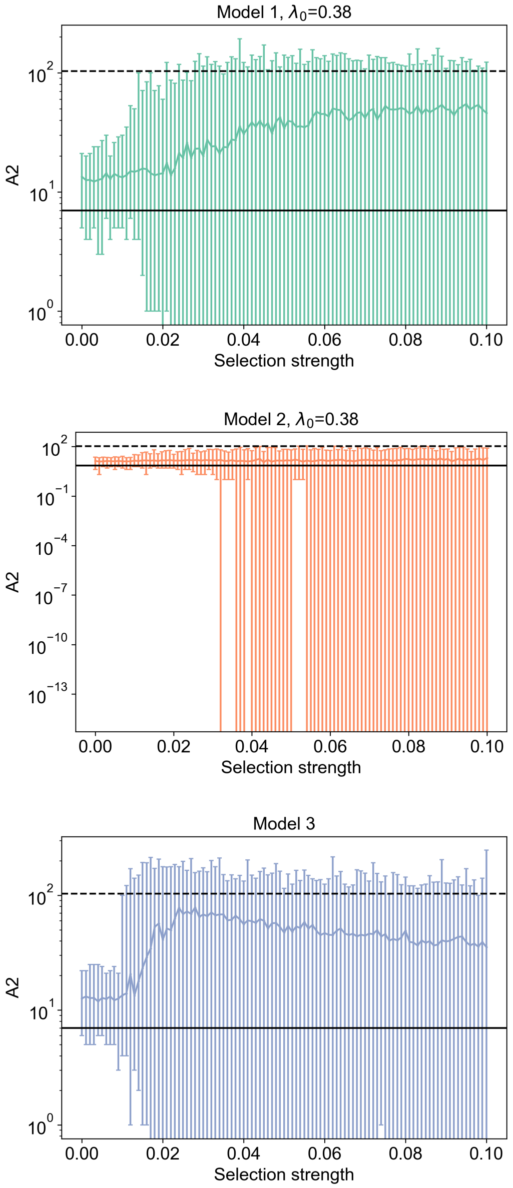


Fig. 16: The variation of the number of truncal mutations detected in all samples (A2) with selection strength $s$. Models 1 and 2 have an assumed death rate of $\lambda_0 = 0.38$. We assume $M = 2$ samples. Here, the absence of a lower dotted line indicates the minimum number of such mutations in tumours from the TRACERx cohort is 0.

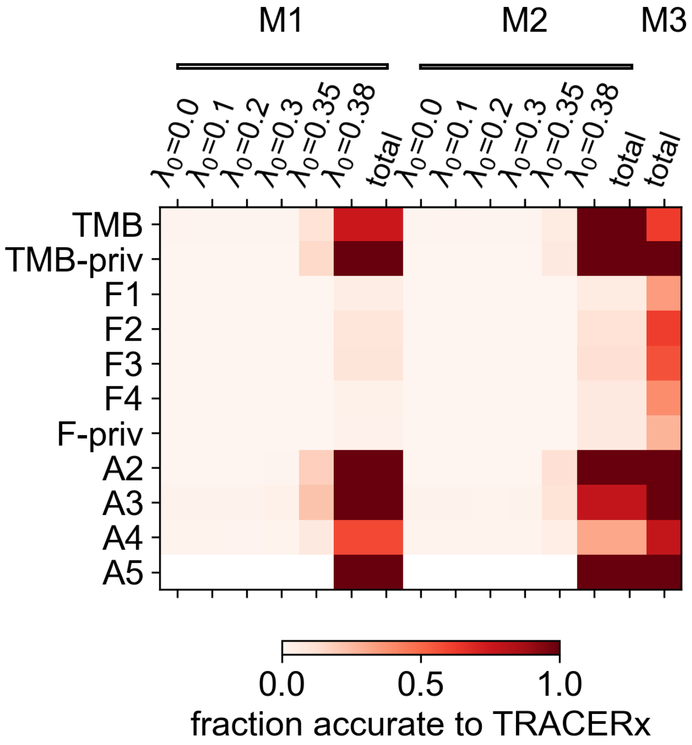


Fig. 17: A comparison of the suitability of all 3 models to the TRACERx data, assuming $M = 5$ samples.

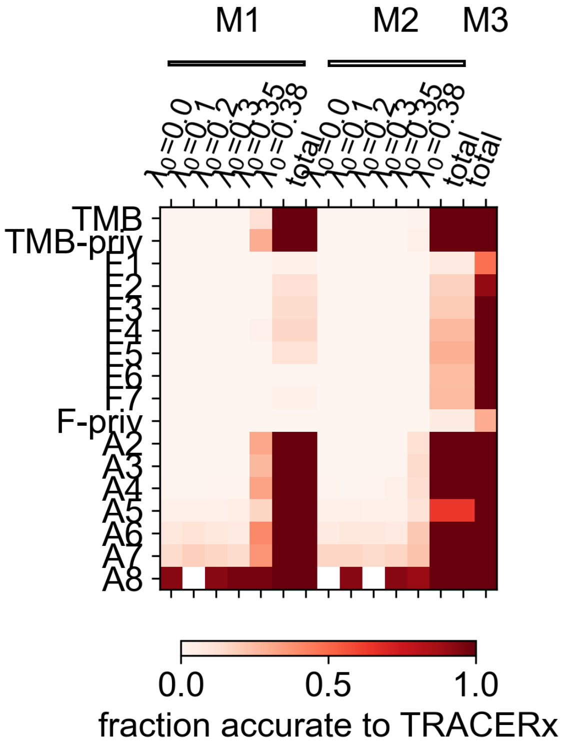


Fig. 18: A comparison of the suitability of all 3 models to the TRACERx data, assuming $M = 8$ samples.

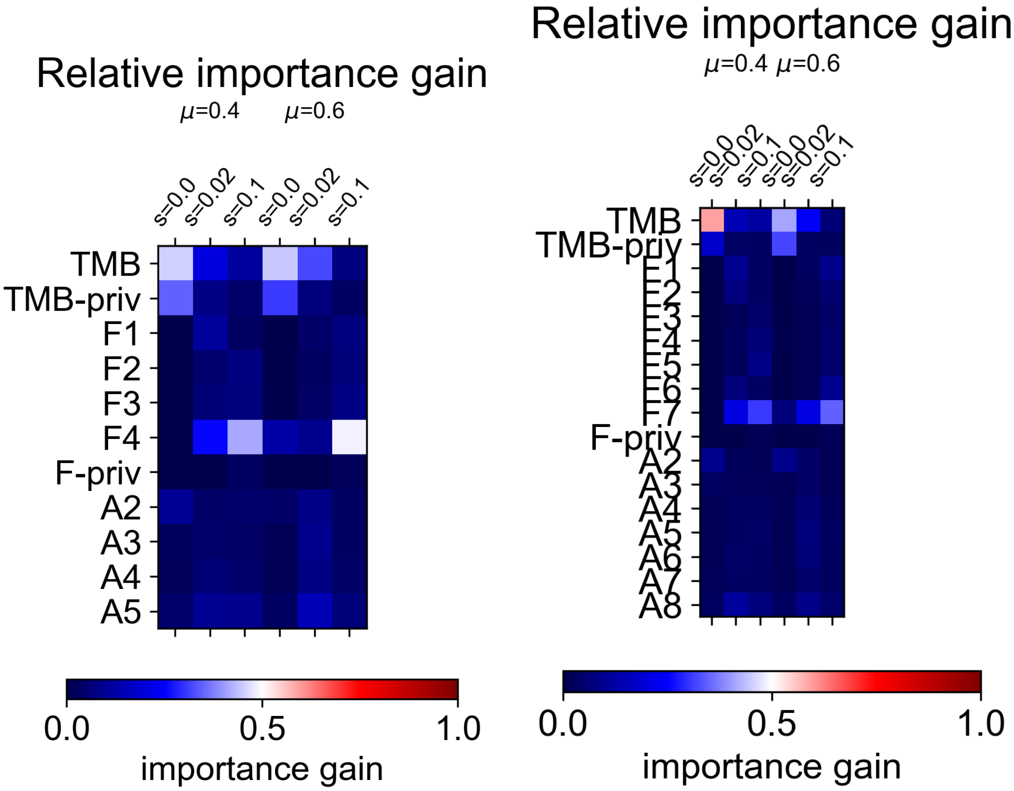


Fig. 19: The relative importance gain for each statistic, assuming $M = 5$ samples (left) or $M = 8$ samples (right). The $(i, j)$th element represents the relative importance gain of summary statistic $i$ to the binary classifier responsible for recognising *in silico* tumours from scenario $j$. Importance gains are normalised across all summary statistics and 10 repeats of the classification process. The objective of XGB-Regressor is to minimise the MSE.

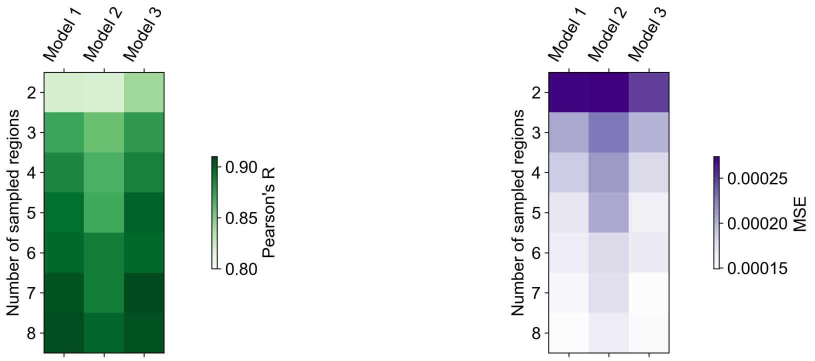


Fig. 20: The accuracy of inference from Models 1, 2, and 3, as measured by the correlation between true and inferred selection strength (Pearson's R, left) or mean squared error (right). Models 1 and 2 have an assumed death rate of $\lambda_0 = 0.38$. The inferred selection is taken as the average of the selection strengths corresponding to the 1% closest tested simulations; error bars indicate the standard deviations of these inferences.

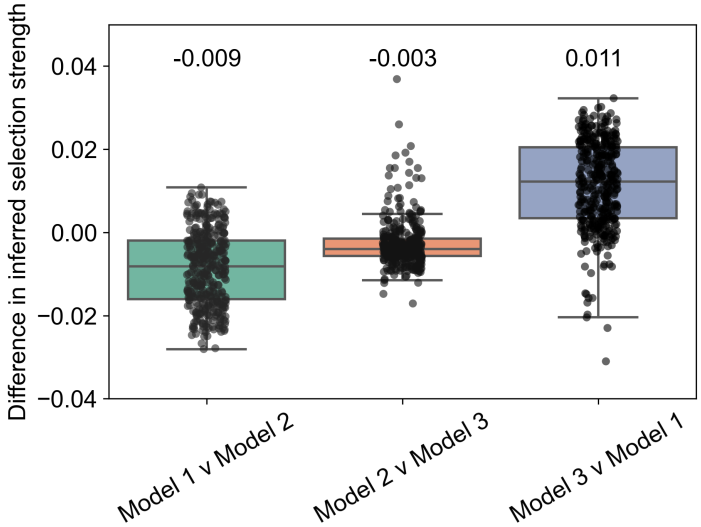


Fig. 21: A box plot of the pairwise differences between model inferences. The $i$th point in leftmost column corresponds to tumour $i$, and represents $s_{i1} - s_{i2}$, where $s_{ij}$ is the average selection strength inferred from tumour $i$ under Model $j$. The average of each distribution is given above each plot. Significance is tested using the distribution $s_{ij} - s_{ik}$, representing the difference in inferred selection levels for tumour $i$ using Models $j$ and $k$; we use a 1-sample T-test with the null-hypothesis that the mean of each distribution is zero, i.e. there is no significant difference between the predictions of Models $j$ and $k$. All three p-values are of order $10^{-30}$ or lower.
